# Conflicts are parametrically encoded: initial evidence for a cognitive space view to reconcile the debate of domain-general and domain-specific cognitive control

**DOI:** 10.1101/2023.02.13.528292

**Authors:** Guochun Yang, Haiyan Wu, Qi Li, Xun Liu, Zhongzheng Fu, Jiefeng Jiang

**Affiliations:** CAS Key Laboratory of Behavioral Science, Institute of Psychology, Beijing 100101, China; Department of Psychology, University of Chinese Academy of Sciences, Beijing 100101, China; Department of Psychological and Brain Sciences, University of Iowa, Iowa City, IA 52242, USA; Cognitive Control Collaborative, University of Iowa, Iowa City, IA 52242, USA; Centre for Cognitive and Brain Sciences and Department of Psychology, University of Macau, Taipa, Macau 999078, China; Beijing Key Laboratory of Learning and Cognition, School of Psychology, Capital Normal University, Beijing 100048, China; Department of Neurosurgery, Cedars-Sinai Medical Center, Los Angeles, CA 90048, USA; Division of Humanities and Social Sciences, California Institute of Technology, Pasadena, CA 91125, USA

**Keywords:** cognitive control, cognitive space, domain-general, domain-specific, conflict

## Abstract

Cognitive control resolves conflicts between task-relevant and -irrelevant information to enable goal-directed behavior. As conflicts can arise from different sources (e.g., sensory input, internal representations), how a limited set of cognitive control processes can effectively address diverse conflicts remains a major challenge. Based on the cognitive space theory, different conflicts can be parameterized and represented as distinct points in a (low-dimensional) cognitive space, which can then be resolved by a limited set of cognitive control processes working along the dimensions. It leads to a hypothesis that conflicts similar in their sources are also represented similarly in the cognitive space. We designed a task with five types of conflicts that could be conceptually parameterized. Both human performance and fMRI activity patterns in the right dorsolateral prefrontal (dlPFC) support that different types of conflicts are organized based on their similarity, thus suggesting cognitive space as a principle for representing conflicts.

## Introduction

Cognitive control enables humans to behave purposefully by modulating neural processing to resolve conflicts between task-relevant and task-irrelevant information. For example, when naming the color of the word “BLUE” printed in red ink, we are likely to be distracted by the word meaning, because reading a word is highly automatic in daily life. To keep our attention on the color, we need to mobilize the cognitive control processes to resolve the conflict between the color and word by boosting/suppressing the processing of color/word meaning. As task-relevant and task-irrelevant information can come from different sources, the sources of conflicts and how they should be resolved can vary greatly (Kornblum et al., 1990). For example, the conflict may occur between items of sensory information, such as between a red light and a police officer signaling cars to pass. Alternatively, conflict may occur between sensory and motor information, such as when a voice on the left asks you to turn right. A key unsolved question in cognitive control is how our brain efficiently resolves these different types of conflicts.

A first step to addressing this question is to examine the commonalities and/or dissociations across different types of conflicts that can be categorized into different *domains*. Examples of the domains of conflicts include experimental paradigm (Freitas et al., 2007; Magen & Cohen, 2007), sensory modality (Hazeltine et al., 2011; Yang et al., 2017), or conflict type regarding the dimensional overlap of conflict processes (Jiang & Egner, 2014; Liu et al., 2004).

Two solutions to resolving different conflict types are proposed. They differ based on whether the same cognitive control mechanisms are applied across domains. On the one hand, the *domain-general* cognitive control theories posit that the frontoparietal cortex adaptively encodes task information and can thus flexibly implement control strategies for different types of conflicts. This is supported by the generalizable control adjustment (i.e., encountering a conflict trial from one type can facilitate conflict resolution of another type) (Freitas et al., 2007; Kan et al., 2013) and similar neural patterns (Peterson et al., 2002; Wu et al., 2020) across distinct conflict tasks. A broader domain-general view holds that the frontoparietal brain regions/networks are widely involved in multiple control demands well beyond the conflict domain (Assem et al., 2020; Cole et al., 2013), which explains the remarkable flexibility in human behaviors. However, since domain-general processes are by definition likely shared by different tasks, when we need to handle multiple task demands at the same time, the efficiency of both tasks would be impaired due to resource competition or interference (Musslick & Cohen, 2021). Therefore, the domain-general processes is evolutionarily less advantageous for humans to deal with the diverse situations requiring high efficiency (Cosmides & Tooby, 1994). On the other hand, the *domain-specific* theories argue that different types of conflicts are handled by distinct cognitive control processes (e.g., where and how information processing should be modulated)(Egner, 2008; Kim et al., 2012). However, according to the domain-specific view, the diverse conflict situations require a multitude of preexisting control processes, which is biologically implausible (Abrahamse et al., 2016).

To reconcile the two theories, researchers recently proposed that cognitive control might be a mixture of domain-general and domain-specific processes. For instance, Freitas and Clark (2015) found that trial-by-trial adjustment of control can generalize across two conflict domains to different degrees, leading to domain-general (strong generalization) or domain-specific (weak or no generalization) conclusions depending on the task settings of the consecutive conflicts. Similarly, different brain networks may show domain-generality (i.e., representing multiple conflicts) or domain-specificity (i.e., representing individual conflicts separately)(Jiang & Egner, 2014; Li et al., 2017). Even within the same brain area (e.g., medial frontal cortex), Fu et al.(2022) found that the neural population activity can be factorized into orthogonal dimensions encoding both domain-general and domain-specific conflict information, which can be selectively read out by downstream brain regions. While the mixture view provides an explanation for the contradictory findings (Braem et al., 2014), it suffers the same criticism as domain-specific cognitive control theories, as it still requires many cognitive control processes to fully cover all possible conflicts.

A key to reconciling domain-general and domain-specific cognitive control is to organize the large number of conflict types using a system with limited, dissociable dimensions. A construct with a similar function is the *cognitive space* (Bellmund et al., 2018), which extends the idea of cognitive map (Behrens et al., 2018) to the representation of abstract information. Critically, the cognitive space view holds that the representations of different abstract information are organized continuously and the representational geometry in the cognitive space is determined by the similarity among the represented information (Bellmund et al., 2018).

In the human brain, it has been shown that abstract (Behrens et al., 2018; Schuck et al., 2016) and social (Park et al., 2020) information can be represented in a cognitive space. For example, social hierarchies with two independent scores (e.g., popularity and competence) can be represented in a 2D cognitive space (one dimension for each score), such that each social item can be located by its score in the two dimensions (Park et al., 2020). In the field of cognitive control, recent studies have begun to conceptualize different control states within a cognitive space (Badre et al., 2021). For example, Fu et al.(2022) mapped different conflict conditions to locations in a low/high dimensional cognitive space to demonstrate the domain-general/domain-specific problems; Grahek et al. (2023) used a cognitive space model of cognitive control settings to explain behavioral changes in the speed-accuracy tradeoff. However, the cognitive spaces proposed in these studies were only applicable to a limited number of control states involved in their designs. Therefore, it remains unclear whether there is a cognitive space that can explain the large number of control states, similar to that of the spatial location (Bellmund et al., 2018) and non-spatial knowledge (Behrens et al., 2018). A challenge to answering this question lies in how to construct control states with continuous levels of similarity. Our recent work (Yang et al., 2021) showed that it is possible to manipulate continuous conflict similarity by using a mixture of two independent conflict types with varying ratios, which can be used to further examine the behavioral and neural evidence for the cognitive space view. It is also unclear how the cognitive space of cognitive control is encoded in the brain, although that of spatial locations and non-spatial abstract knowledge has been relatively well investigated in the medial temporal lobe, medial prefrontal and orbitofrontal system (Behrens et al., 2018; Bellmund et al., 2018). Recent research has suggested that the abstract task structure could be encoded and implemented by the frontoparietal network (Vaidya & Badre, 2022; Vaidya et al., 2021), but whether a similar neural system encodes the cognitive space of cognitive control remains untested.

The present study aimed to test the geometry of cognitive space in conflict representation. Specifically, we hypothesize that different types of conflicts are represented as points in a cognitive space. Importantly, the distance between the points, which reflects the geometry of the cognitive space, scales with the difference in the sources of the conflicts being represented by the points. The dimensions in the cognitive space of conflicts can be the aforementioned *domains*, in which domain-specific cognitive control processes are defined. For a specific type of conflict, its location in the cognitive space can be parameterized using a limited number of coordinates, which reflect how much control is needed for each of the domain-specific cognitive control processes. The cognitive space can also represent different types of conflicts with low dimensionality (Badre et al., 2021; MacDowell et al., 2022). Different domains can be represented conjunctively in a single cognitive space to achieve domain-general cognitive control, as conflicts from different sources can be resolved using the same set of cognitive control processes. We further hypothesize that the cognitive space representing different types of conflicts may be located in the frontoparietal network due to its essential roles in conflict resolution (Freund, Bugg, et al., 2021; Fu et al., 2022) and abstract task representation (Vaidya & Badre, 2022).

In this study, we adjusted the paradigm from our previous study (Yang et al., 2021) by including transitions of trials from five different conflict types, which enabled us to test if these conflict types are organized in a cognitive space (Fig. 1A). Specifically, on each trial, an arrow, pointing either upwards or downwards, was presented on one of the 10 possible locations on the screen. Participants were required to respond to the pointing direction of the arrow (up or down) by pressing either the left or right key. Importantly, conflicts from two sources can occur in this task. On one hand, the vertical location of the arrow can be incongruent with the direction (e.g., an up-pointing arrow on the lower half of the screen), resulting spatial Stroop conflict (Liu et al., 2004; Lu & Proctor, 1995). On the other hand, the horizontal location of the arrow can be incongruent with the response key (e.g., an arrow requiring left response presented on the right side of the screen), thus causing Simon conflict (Lu & Proctor, 1995; Simon & Small, 1969). As the arrow location rotates from the horizontal axis to the vertical axis, spatial Stroop conflict increases, and Simon conflict decreases. Therefore, the 10 possible locations of the arrow give rise to five conflict types with unique blend of spatial Stroop and Simon conflicts (Yang et al., 2021). As the increase in spatial Stroop conflict is highly correlated with the decrease in Simon conflict, we can use a 1D cognitive space to represent all five conflict types.

**Fig. 1.**
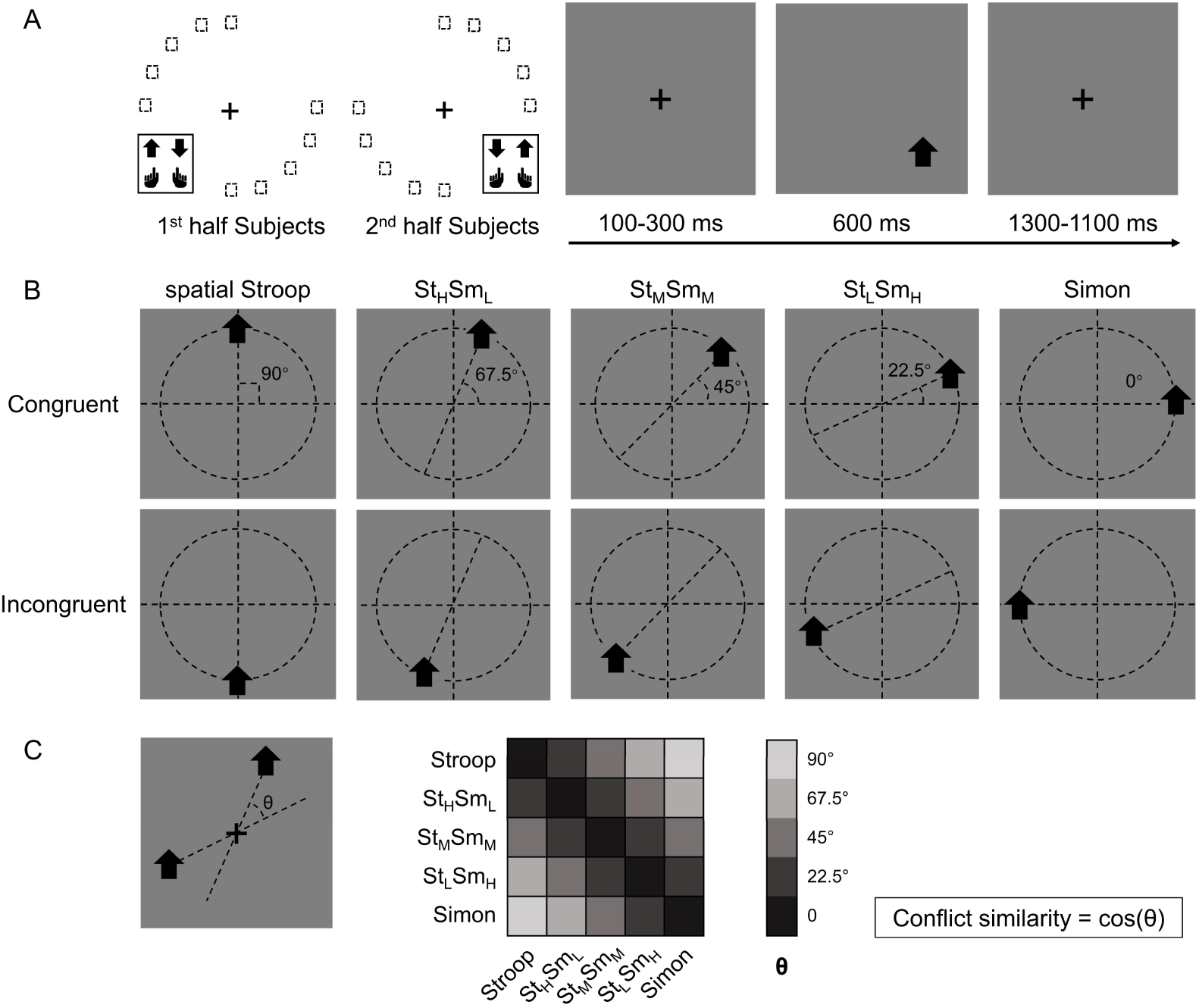
Experimental design. (A) The left panel shows the orthogonal stimulus-response mappings of the two participant groups. In each group the stimuli were only displayed at two quadrants of the circular locations. One group were asked to respond with the left button to the upward arrow and with the right button to the downward arrow presented in the top-left and bottom-right quadrants, and the other group vice versa. The right panel shows the time course of one example trial. The stimuli were displayed for 600 ms, preceded and followed by fixation crosses that lasted for 1400 ms in total. (B) Examples of the five types of conflicts, each containing congruent and incongruent conditions. The arrows were presented at locations along five orientations with isometric polar angles, in which the vertical location introduces the spatial Stroop conflict, and the horizontal location introduces the Simon conflict. Dashed lines are shown only to indicate the location of arrows and were not shown in the experiments. (C) The definition of the angular difference between two conflict types and the conflict similarity. The angle θ is determined by the acute angle between two lines that cross the stimuli and the central fixation. Therefore, stimuli of the same conflict type form the smallest angle of 0, and stimuli between Stroop and Simon form the largest angle of 90°, and others are in between. Conflict similarity is defined by the cosine value of θ. H = high; L = low; M = medium.

One way to parameterize (i.e., defining a coordinate system) the cognitive space is to encode each conflict type by the angle of the axis connecting its two possible stimulus locations (Fig. 1B). Within this cognitive space, the similarity between two conflict types can be quantified as the cosine value of their angular difference (Fig. 1C). The rationale behind defining conflict similarity based on combinations of different conflict sources, such as spatial-Stroop and Simon, stems from the evidence that these sources undergo independent processing (Egner, 2008; Li et al., 2014; Liu et al., 2010; Wang et al., 2014). Identifying these distinct sources is critical in efficiently resolving diverse conflicts. If the conflict types are organized as a cognitive space in the brain, the similarity between conflict types in the cognitive space should be reflected in both the behavior and similarity in the neural representations of conflict types. Our data from two experiments using this experimental design support both predictions: using behavioral data, we found that the influence of congruency (i.e., whether the task-relevant and task-irrelevant information indicate the same response) from the previous trial to the next trial increases with the conflict similarity between the two trials. Using fMRI data, we found that more similar conflicts showed higher multivariate pattern similarity in the right dorsolateral prefrontal cortex (dlPFC).

## Results

### Conflict type similarity modulates behavioral congruency sequence effect (CSE)

#### Experiment 1

We conducted a behavioral experiment (n = 33, 18 females) to examine how CSEs across different conflict types are influenced by their similarity. First, we validated the experimental design by testing the congruency effects. All five conflict types showed robust congruency effects such that the incongruent trials were slower and less accurate than the congruent trials (Note S1; Fig. S1 A/B). To test the influence of similarity between conflict types on behavior, we examined the CSE in consecutive trials. Specifically, the CSE was quantified as the interaction between previous and current trial congruency and can reflect how (in)congruency on the previous trial influences cognitive control on the current trial (Egner, 2007; Schmidt & Weissman, 2014). It has been shown that the CSE diminishes if the two consecutive trials have different conflict types (Akcay & Hazeltine, 2011; Egner et al., 2007; Torres-Quesada et al., 2013). Similarly, we tested whether the size of CSE increases as a function of conflict similarity between consecutive trials. To this end, we organized trials based on a 5 (previous trial conflict type) × 5 (current trial conflict type) × 2 (previous trial congruency) × 2 (current trial congruency) factorial design, with the first two and the last two factors capturing between-trial conflict similarity and the CSE, respectively. The cells in the 5 × 5 matrix were mapped to different similarity levels according to the angular difference between the two conflict types (Fig. 1C). As shown in Fig. 2, the CSE, measured in both reaction time (RT) and error rate (ER), scaled with conflict similarity.

**Fig. 2.**
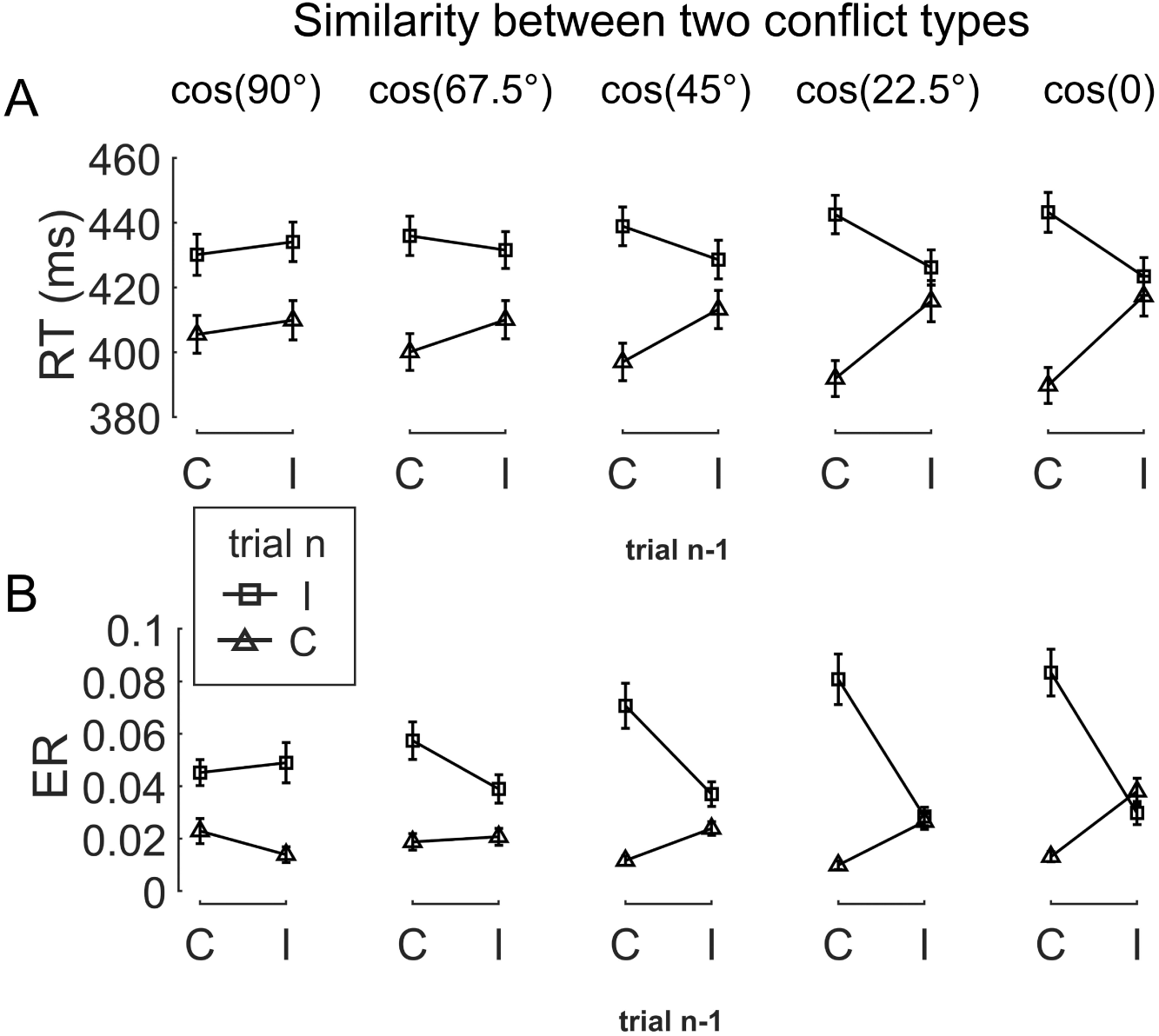
The conflict similarity modulation on the behavioral CSE in Experiment 1. (A) RT and (B) ER are plotted as a function of congruency types on trial n−1 and trial n. Each column shows one similarity level, as indicated by the defined angular difference between two conflict types. Error bars are standard errors. C = congruent; I = incongruent; RT = reaction time; ER = error rate.

To test the modulation of conflict similarity on the CSE, we constructed a linear mixed effect model to predict RT/ER in each cell of the factorial design using a predictor encoding the interaction between the CSE and conflict similarity (see Methods). The results showed a significant effect of conflict similarity (RT: *β =* 0.10 ± 0.01, *t*(1719.7) = 15.82, *p* < .001, *η_p_*^2^ = .120; ER: *β =* 0.15 ± 0.02, *t*(204.5) = 7.84, *p* < .001, *η_p_*^2^ = .085, Fig. S2B/E). In other words, the CSE increased with the conflict similarity between two consecutive trials. The conflict similarity modulation effect remained significant after regressing out the influence of physical proximity between the stimuli of consecutive trials (Note S2). As a control analysis, we also compared this approach to a two-stage analysis that first calculated the CSE for each previous × current trial conflict type condition and then tested the modulation of conflict similarity on the CSEs (Yang et al., 2021). The two-stage analysis also showed a strong effect of conflict similarity (RT: *β =* 0.58 ± 0.04, *t*(67.5) = 14.74, *p* < .001, *η_p_*^2^ = .388; ER: *β =* 0.36 ± 0.05, *t*(40.3) = 7.01, *p* < .001, *η_p_*^2^ = .320, Fig. S2A/D). Importantly, individual modulation effects of conflict similarity were positively correlated between the two approaches (RT: *r =* 0.48; ER: *r =* 0.86, both *p*s < 0.003, one-tailed), indicating the consistency of the estimated conflict similarity effects across the two approaches. In the following texts, we will use the terms “*conflict similarity effect”* and “*conflict type effect*” interchangeably.

Moreover, to test the continuity and generalizability of the similarity modulation, we conducted a leave-one-out prediction analysis. We used the behavioral data from Experiment 1 for this test, due to its larger amount of data than Experiment 2. Specifically, we removed data from one of the five similarity levels (as illustrated by the *θ*s in Fig. 1C) and used the remaining data to perform the same mixed-effect model (i.e., the two-stage analysis). This yielded one pair of beta coefficients including the similarity regressor and the intercept for each subject, with which we predicted the CSE for the removed similarity level for each subject. We repeated this process for each similarity level once. The predicted results were highly correlated with the original data, with *r* = .87 for the RT and *r* = .84 for the ER, *p*s < .001.

#### Experiment 2

##### Behavioral results

We next conducted an fMRI experiment using a shorter version of the same task with a different sample (n = 35, 17 females) to seek neural evidence of how different conflict types are organized. Using behavioral data, we first validated the experimental design by testing congruency effects in each of the five conflict types (Note S1; Fig. S1 C/D). We then tested the modulation of conflict similarity on the behavioral CSE using the linear mixed effect model as in Experiment 1 (except the two-stage method). Results showed a significant effect of conflict similarity modulation (RT: *β =* 0.24 ± 0.04, *t*(71.7) = 6.36, *p* < .001, *η_p_*^2^ = .096; ER: *β =* 0.33 ± 0.06, *t*(175.4) = 5.81, *p* < .001, *η_p_*^2^ = .124, Fig. S2C/F), thus replicating the results of Experimental 1 and setting the stage for fMRI analysis. As in Experiment 1, the conflict similarity modulation effect remained significant after regressing out the influence of physical proximity between the stimuli of consecutive trials (Note S2).

### Univariate brain activations scale with conflict strength

In the fMRI analysis, we first replicated the classic congruency effect by searching for brain regions showing higher univariate activation in incongruent than congruent conditions (GLM1, see Methods). Consistent with the literature (Botvinick et al., 2004; Fu et al., 2022), this effect was observed in the pre-supplementary motor area (pre-SMA) (Fig. 3, Table S1). We then tested the encoding of conflict type as a cognitive space by identifying brain regions with activation levels parametrically covarying with the coordinates (i.e., axial angle relative to the horizontal/vertical axes) in the hypothesized cognitive space. As shown in Fig. 1B, change in the angle corresponds to change in spatial Stroop and Simon conflicts in opposite directions, so we used opposite contrasts to examine the encoding of spatial Stroop and Simon strength, respectively (see Methods). Accordingly, we found the right inferior parietal sulcus (IPS) and the right dorsomedial prefrontal cortex (dmPFC) displayed positive correlation between fMRI activation and the Simon conflict (Fig. 3, Fig. S3, Table S1). We did not observe regions showing significant correlation with the spatial Stroop conflict.

**Fig. 3.**
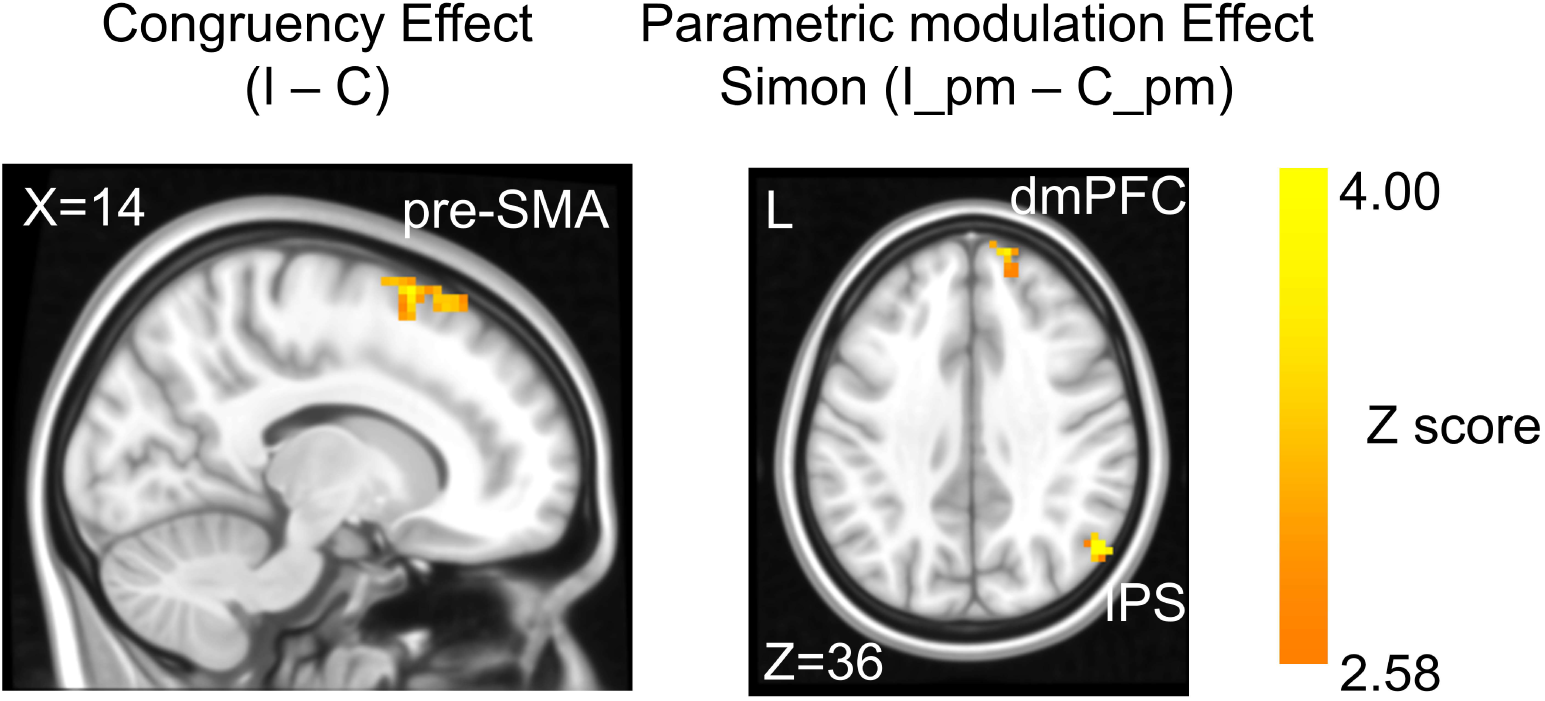
The congruency effect and parametric modulation effect detected by uni-voxel analyses. Results displayed are probabilistic TFCE enhanced and thresholded with voxel-wise *p* < .001 and cluster-wise *p* < .05, both one-tailed. The congruency effect denotes the higher activation in incongruent than congruent condition (left panel). The positive parametric modulation effect (I_pm – C_pm) denotes the higher activation when the conflict type contained a higher ratio of Simon conflict component (right panel). I = incongruent; C = congruent; pm = parametric modulator.

To further test if the univariate results explain the conflict similarity modulation of the behavioral CSE (slope in Fig. S2C), we conducted brain-behavioral correlation analyses for regions identified above. Regions with higher spatial Stroop/Simon modulation effects were expected to trigger higher behavioral conflict similarity modulation effect on the CSE. However, none of the two regions (i.e., right IPS and right dmPFC, Fig. 3) were positively correlated with the behavioral performance, both uncorrected *ps* >.28, one-tailed. In addition, since the conflict type difference covaries with the orientation of the arrow location at the individual level (e.g., the stimulus in a higher level of Simon conflict is always closer to the horizontal axis, see Fig. 4A), the univariate modulation effects may not reflect purely conflict type difference. To further tease these factors apart, we used multivariate analyses.

**Fig. 4.**
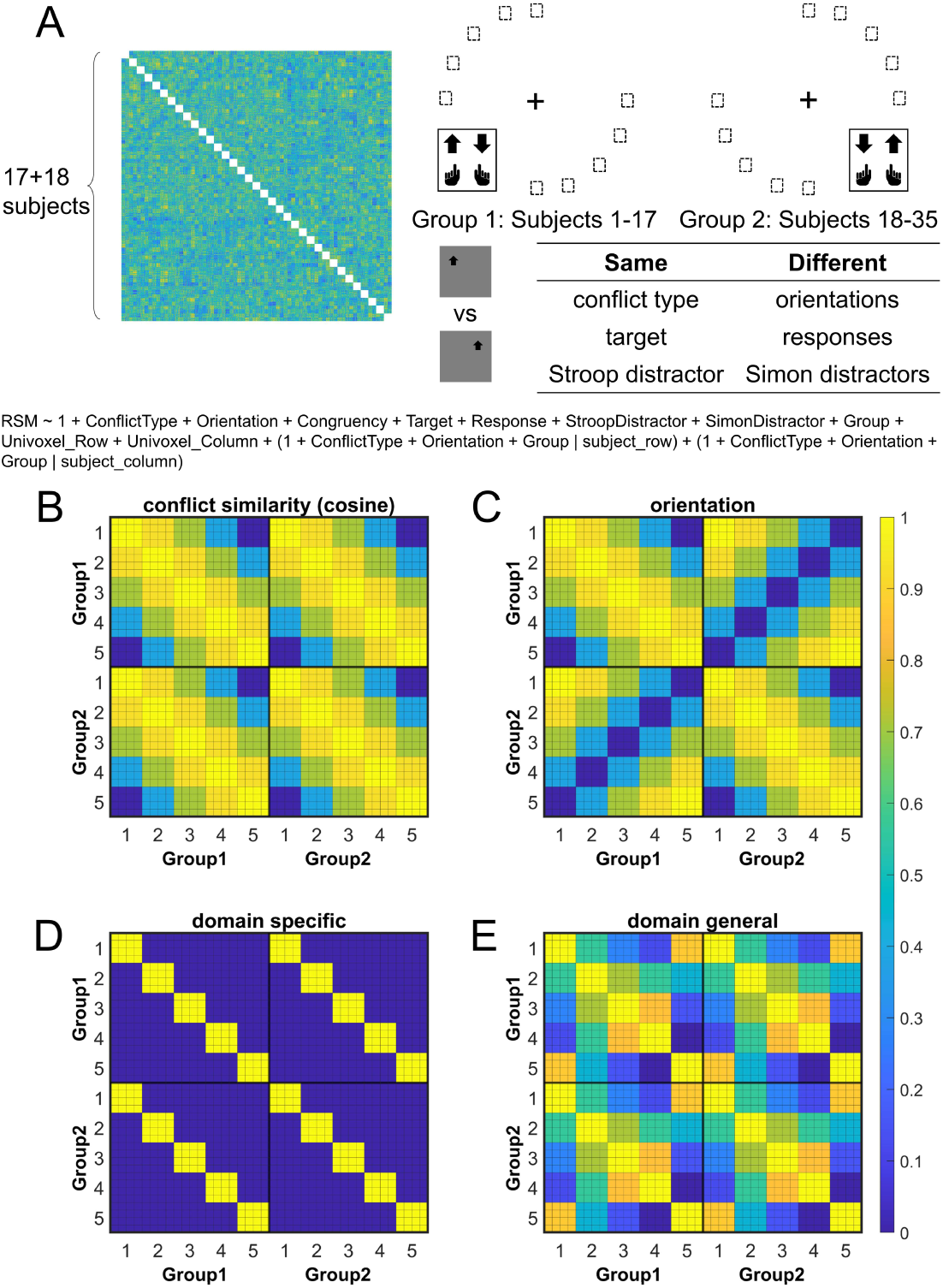
Rationale of the cross-subject RSA model and the schematic of key RSMs. A) The RSM is calculated as the Pearson’s correlation between each pair of conditions across the 35 subjects. For 17 subjects, the stimuli were displayed on the top-left and bottom-right quadrants, and they were asked to respond with left hand to the upward arrow and right hand to the downward arrow. For the other 18 subjects, the stimuli were displayed on the top-right and bottom-left quadrants, and they were asked to respond with left hand to the downward arrow and right hand to the upward arrow. Within each subject, the conflict type and orientation regressors were perfectly covaried. For instance, the same conflict type will always be on the same orientation. To de-correlate conflict type and orientation effects, we conducted the RSA across subjects from different groups. For example, the bottom-right panel highlights the example conditions that are orthogonal to each other on the orientation, response, and Simon distractor, whereas their conflict type, target and spatial Stroop distractor are the same. The dashed boxes show the possible target locations for different conditions. (B) and (C) show the orthogonality between conflict similarity and orientation RSMs. The within-subject RSMs (e.g., Group1-Group1) for conflict similarity and orientation are all the same, but the cross-group correlations (e.g., Group2-Group1) are different. Therefore, we can separate the contribution of these two effects when including them as different regressors in the same linear regression model. (D) and (E) show the two alternative models. Like the cosine model (B), within-group trial pairs resemble between-group trial pairs in these two models. The domain-specific model is an identity matrix. The domain-general model is estimated from the absolute difference of behavioral congruency effect, but scaled to 0 (lowest similarity) – 1 (highest similarity) to aid comparison. The plotted matrices in B-E include only one subject each from Group 1 and Group 2. Numbers 1-5 indicate the conflict type conditions, for spatial Stroop, St_H_Sm_L_, St_M_Sm_M_, St_L_Sm_H_, and Simon, respectively. The thin lines separate four different sub-conditions, i.e., target arrow (up, down) × congruency (incongruent, congruent), within each conflict type.

### Multivariate patterns of the right dlPFC encodes the conflict similarity

The hypothesis that the brain encodes conflict types in a cognitive space predicts that similar conflict types will have similar neural representations. To test this prediction, we computed the representational similarity matrix (RSM) that encoded correlations of blood-oxygen-level dependent (BOLD) signal patterns between each pair of conflict type (Stroop, St_H_Sm_L_, St_M_Sm_M_, St_L_Sm_H_, and Simon, with H, M and L indicating high, medium and low, respectively, see also Fig. 1B) × congruency (congruent, incongruent) × arrow direction (up, down) × run × subject combinations for each of the 360 cortical regions from the Multi-Modal Parcellation (MMP) cortical atlas (Glasser et al., 2016; Jiang et al., 2020). The RSM was then submitted to a linear mixed-effect model as the dependent variable to test whether the representational similarity in each region was modulated by various experimental variables (e.g., conflict type, spatial orientation, stimulus, response, etc., see Methods). The linear mixed-effect model was used to de-correlate conflict type and spatial orientation leveraging the between-subject manipulation of stimulus locations (Fig. 4A).

To validate this method, we applied this analysis to test the effects of response/stimulus features and found that representational similarity of the BOLD signal patterns significantly covaried with whether two response/spatial location/arrow directions were the same most strongly in bilateral motor/visual/somatosensory areas, respectively (Fig. S5). We then identified the cortical regions encoding conflict type as a cognitive space by testing whether their RSMs can be explained by the similarity between conflict types. Specifically, we applied three independent criteria: (1) the cortical regions should exhibit a statistically significant positive conflict similarity effect on the RSM; (2) the conflict similarity effect should be stronger in incongruent than congruent trials to reflect flexible adjustment of cognitive control demand when the conflict is present; and (3) the conflict similarity effect should be positively correlated with the behavioral conflict similarity modulation effect on the CSE (see *Behavioral Results* of Experiment 2). The first criterion revealed several cortical regions encoding the conflict similarity, including the frontal eye field (FEF), region 1, Brodmann 8C area (a subregion of dlPFC) (Glasser et al., 2016), a47r, posterior inferior frontal junction (IFJp), anterior intraparietal area (AIP), temporoparietooccipital junction 3 (TPOJ3), PGi, and V3CD in the right hemisphere, and the superior frontal language (SFL) area, 23c, 24dd, 7Am, p32pr, 6r, FOP1, PF, ventromedial visual area (VMV1/2) areas, area 25, MBelt in the left hemisphere (Bonferroni corrected *ps* < 0.05, one-tailed, Fig. 5A). We next tested whether these regions were related to cognitive control by comparing the strength of conflict similarity effect between incongruent and congruent conditions (criterion 2) and correlating the strength to behavioral similarity modulation effect (criterion 3). Given these two criteria pertain to second-order analyses (interaction or individual analyses) and thus might have lower statistical power (Blake & Gangestad, 2020), we applied a more lenient threshold using false discovery rate (FDR) correction (Benjamini & Hochberg, 1995) on the above-mentioned regions. Results revealed that the left SFL, left VMV1, area 1eft 25 and right 8C met this criterion, FDR corrected *p*s < .05, one-tailed, suggesting that the representation of conflict type was strengthened when the conflict was present (e.g., Fig. 5D and Fig. S4). The inter-subject brain-behavioral correlation analysis (criterion 3) showed that the strength of conflict similarity effect on RSM scaled with the modulation of conflict similarity on the CSE (slope in Fig. S2C) in right 8C (*r* = .52, FDR corrected *p* = .015, one-tailed, Fig. 5C) only. These results are listed in Table 1. In addition, we did not find evidence supporting the encoding of congruency in the right 8C area (see Note S6), suggesting that the right 8C area specifically represents conflict similarity.

**Fig. 5.**
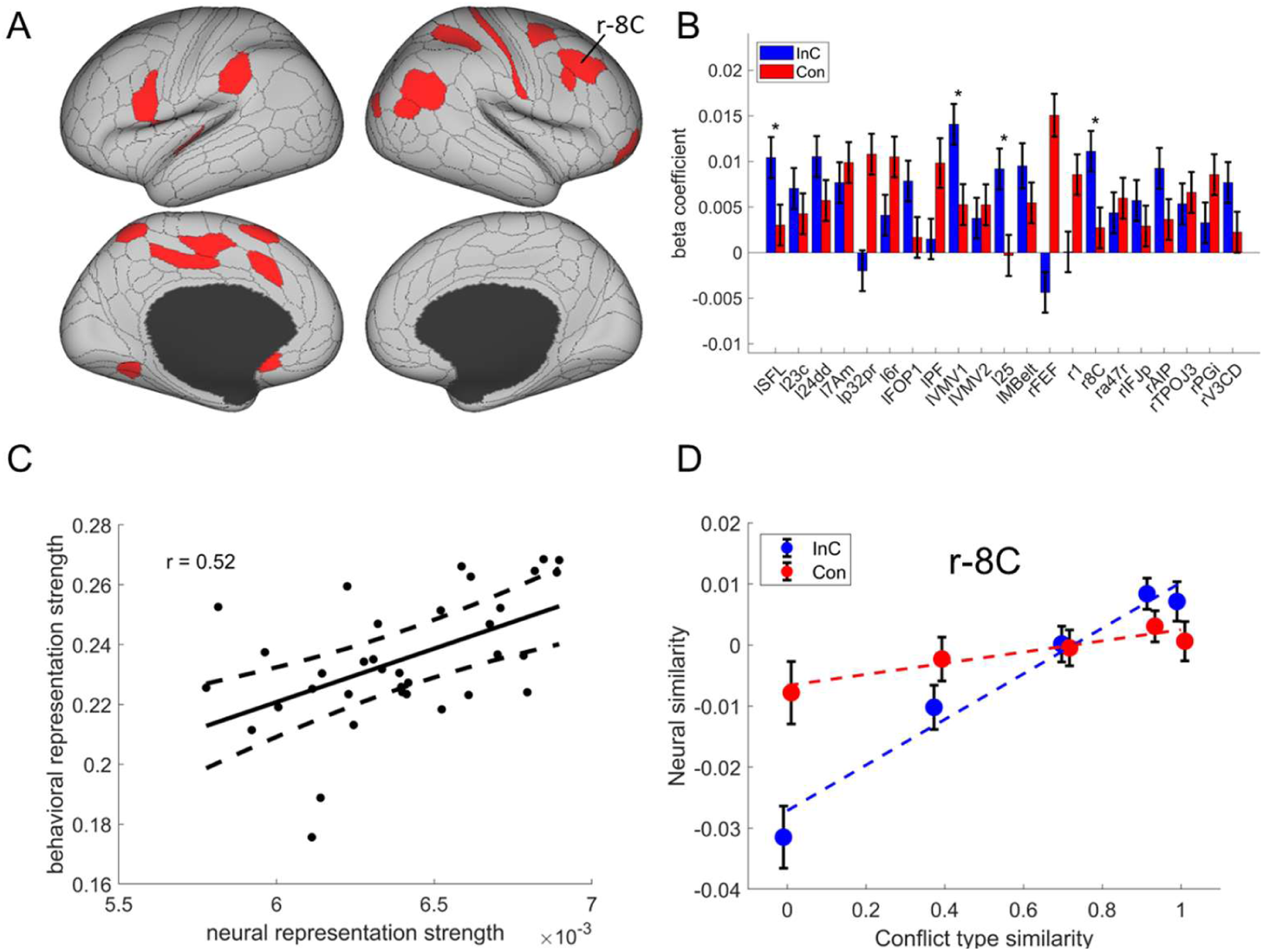
The conflict type effect. (A) Brain regions surviving the Bonferroni correction (*p* < 0.05) across the 360 regions (criterion 1). Labeled is the region meeting the three criteria. (B) Different encoding of conflict type effect in the incongruent with congruent conditions (criterion 2). * FDR corrected *p* < .05; r = right. (C) The brain-behavior correlation of the right 8C (criterion 3). The x-axis shows the beta coefficient of the conflict type effect from the RSA, and the y-axis shows the beta coefficient obtained from the behavioral linear model using the conflict similarity to predict the CSE in Experiment 2. (D) Illustration of the different encoding strength of conflict type similarity in incongruent versus congruent conditions of right 8C. The y-axis is derived from the z-scored Pearson correlation coefficient after regressing out other factors. See Fig. S4B for a plot with the raw Pearson correlation measurement. l = left; r = right.

**Table 1.** Summary statistics of the cross-subject RSA for regions showing conflict type and orientation effects identified by the three criteria.

| Region<br>name | Criterion 1 |  |  | Criterion 2 |  | Criterion 3 |  |
| --- | --- | --- | --- | --- | --- | --- | --- |
| | *t* | $\beta \pm \text{SD}$ | *p* | *t* | *p* | *r* | *p* |
| *Conflict type effect* |  |  |  |  |  |  |  |
| left SFL | 4.77 | 0.0061 ± 0.0013 | 1.9$\times 10^{-6}$ | 2.45 | 0.007 | .35 | .021 |
| left 23c | 4.42 | 0.0049 ± 0.0011 | 9.8$\times 10^{-6}$ | 1.18 | 0.118 | -.15 | .800 |
| left 24dd | 6.13 | 0.0079 ± 0.0013 | 8.9$\times 10^{-10}$ | 1.61 | 0.053 | -.04 | .580 |
| left 7Am | 6.76 | 0.0090 ± 0.0013 | 1.4$\times 10^{-11}$ | -0.75 | 0.772 | .04 | .418 |
| left p32pr | 4.00 | 0.0044 ± 0.0011 | 6.4$\times 10^{-5}$ | -3.95 | 1.000 | -.01 | .533 |
| left 6r | 5.41 | 0.0071 ± 0.0013 | 6.3$\times 10^{-8}$ | -1.94 | 0.973 | -.11 | .737 |
| left FOP1 | 4.37 | 0.0050 ± 0.0011 | 1.3$\times 10^{-5}$ | 1.83 | 0.034 | -.25 | .926 |
| left PF | 4.04 | 0.0058 ± 0.0014 | 5.3$\times 10^{-5}$ | -1.90 | 0.969 | -.24 | .921 |
| left VMV1 | 7.45 | 0.0091 ± 0.0012 | 9.6$\times 10^{-14}$ | 2.83 | 0.002 | -.27 | .940 |
| left VMV2 | 4.75 | 0.0053 ± 0.0011 | 2.0$\times 10^{-6}$ | -0.32 | 0.624 | -.06 | .630 |
| left 25 | 3.70 | 0.0041 ± 0.0011 | 2.1$\times 10^{-4}$ | 3.26 | 0.001 | .05 | .386 |
| left Mbelt | 4.25 | 0.0064 ± 0.0015 | 2.1$\times 10^{-5}$ | 1.77 | 0.039 | .10 | .275 |
| right FEF | 3.98 | 0.0054 ± 0.0014 | 6.8$\times 10^{-5}$ | -5.49 | 1.000 | .08 | .327 |
| right 1 | 3.90 | 0.0045 ± 0.0012 | 9.7$\times 10^{-5}$ | -2.34 | 0.990 | -.07 | .665 |
| right 8C* | 5.41 | 0.0064 ± 0.0012 | 6.1$\times 10^{-8}$ | 2.46 | 0.007 | .52 | .001 |
| right a47r | 5.04 | 0.0056 ± 0.0011 | 4.7$\times 10^{-7}$ | -0.68 | 0.753 | .05 | .393 |
| right IFJp | 3.78 | 0.0042 ± 0.0011 | 1.6$\times 10^{-4}$ | 0.77 | 0.221 | .27 | .056 |
| right AIP | 4.64 | 0.0054 ± 0.0012 | 3.5$\times 10^{-6}$ | 1.86 | 0.032 | -.02 | .540 |
| right TPOJ3 | 4.48 | 0.0056 ± 0.0012 | 7.6$\times 10^{-6}$ | -0.25 | 0.600 | .21 | .118 |
| right PGi | 4.07 | 0.0045 ± 0.0011 | 4.8$\times 10^{-5}$ | -1.59 | 0.944 | -.01 | .523 |
| right V3CD | 3.86 | 0.0043 ± 0.0011 | 1.2$\times 10^{-4}$ | 1.94 | 0.026 | .02 | .451 |
| *Orientation effect* |  |  |  |  |  |  |  |
| left FEF* | 5.17 | 0.0060 ± 0.0012 | 2.4$\times 10^{-7}$ | 2.73 | 0.003 | -.01 | .518 |
| left POS1 | 4.35 | 0.0051 ± 0.0012 | 1.4$\times 10^{-5}$ | -1.52 | 0.936 | .00 | .500 |
| left 31pv | 5.60 | 0.0103 ± 0.0018 | 2.1$\times 10^{-8}$ | -0.48 | 0.686 | .05 | .397 |
| left 6ma | 4.75 | 0.0055 ± 0.0012 | 2.0$\times 10^{-6}$ | -1.87 | 0.970 | -.00 | .500 |
| left 7PC | 3.66 | 0.0042 ± 0.0012 | 2.5$\times 10^{-4}$ | 0.76 | 0.223 | -.20 | .876 |
| left 8BL | 4.55 | 0.0053 ± 0.0012 | 5.3$\times 10^{-6}$ | 0.10 | 0.460 | .14 | .207 |
| left AIP | 3.83 | 0.0044 ± 0.0012 | 1.3$\times 10^{-4}$ | 0.62 | 0.266 | -.23 | .910 |
| left TE1p | 3.95 | 0.0046 ± 0.0012 | 7.8$\times 10^{-5}$ | 0.14 | 0.443 | .01 | .475 |
| left IP2* | 4.49 | 0.0052 ± 0.0012 | 7.2$\times 10^{-6}$ | 4.02 | 0.000 | .19 | .139 |
| right V1* | 4.71 | 0.0083 ± 0.0018 | 2.5$\times 10^{-6}$ | 2.91 | 0.002 | -.08 | .672 |
| right V2* | 3.88 | 0.0172 ± 0.0044 | 1.1$\times 10^{-4}$ | 3.19 | 0.001 | .02 | .462 |
| right V3 | 3.94 | 0.0175 ± 0.0045 | 8.2$\times 10^{-5}$ | -1.45 | 0.927 | .39 | .010 |
| right LO2 | 5.95 | 0.0069 ± 0.0012 | 2.8$\times 10^{-9}$ | -1.81 | 0.965 | .26 | .068 |
| right POS1* | 3.70 | 0.0043 ± 0.0012 | 2.1$\times 10^{-4}$ | 2.98 | 0.001 | .33 | .028 |
| right 5m | 5.24 | 0.0061 ± 0.0012 | 1.6×10<sup>-7</sup> | -2.12 | 0.983 | .00 | .500 |
| right TF | 4.97 | 0.0058 ± 0.0012 | 6.7×10<sup>-7</sup> | -1.08 | 0.860 | -.07 | .657 |
| right PHT | 4.54 | 0.0053 ± 0.0012 | 5.5×10<sup>-6</sup> | 0.03 | 0.486 | -.04 | .589 |
| right PF* | 5.57 | 0.0064 ± 0.0012 | 2.6×10<sup>-8</sup> | 3.08 | 0.001 | -.03 | .558 |
| right A4 | 4.11 | 0.0048 ± 0.0012 | 3.9×10<sup>-5</sup> | -2.68 | 0.996 | -.09 | .700 |

Note. all *p* values listed are 1-tailed and uncorrected. The \* denotes the regions meeting all three criteria for each effect.

The model described above employs the cosine similarity measure to define conflict similarity and will be referred to as the Cognitive-Space model. To examine if the right 8C specifically encodes the cognitive space rather than the domain-general or domain-specific organizations, we tested two additional models (see Methods). Model comparison showed a lower BIC in the Cognitive-Space model (BIC = 5377093) than the Domain-General (BIC = 537126) or Domain-Specific (BIC = 537127) models. Further analysis showed the dimensionality of the representation in the right 8C was 1.19, suggesting the cognitive space was close to 1D. Moreover, we replicated the results with only incongruent trials, considering that the pattern of conflict representations is more manifested when the conflict is present (i.e., on incongruent trials) than not (i.e., on congruent trials). We found a poorer fitting in Domain-general (BIC = 1344127), and Domain-Specific (BIC = 1344129) models than the Cognitive-Space model (BIC = 1344104). These results indicate that the right 8C encodes an integrated cognitive space for resolving Stroop and Simon conflicts. The more detailed model comparison results are listed in Table 2.

**Table 2.** Model comparison results of the right 8C. RSM_I shows results using incongruent trials only.

| Model name | Full RSM |  |  | RSM_I |  |  |
| --- | --- | --- | --- | --- | --- | --- |
|  | *t* | *p* | BIC | *t* | *p* | BIC |
| Cognitive-Space | 5.41 | 6.1×10<sup>-8</sup> | 5377093 | 4.97 | 3.35×10<sup>-7</sup> | 1345201 |
| Domain-General | 0.92 | .179 | 5377126 | 1.43 | .076 | 1344127 |
| Domain-Specific | 0.84 | .200 | 5377127 | 0.28 | .390 | 1344129 |

In sum, we found converging evidence supporting that the right dlPFC (8C area) encoded conflict similarity parametrically, which further supports the hypothesis that conflict types are represented in a cognitive space.

### Multivariate patterns of visual and oculomotor areas encode stimulus orientation

To tease apart the representation of conflict type from that of perceptual information, we tested the modulation of the spatial orientations of stimulus locations on RSM using the aforementioned RSA. We also applied three independent criteria: (1) the cortical regions should exhibit a statistically significant orientation effect on the RSM; (2) the conflict similarity effect should be stronger in incongruent than congruent trials; and (3) the orientation effect should not interact with the CSE, since the orientation effect was dissociated from the conflict similarity effect and was not expected to influence cognitive control. We observed increasing fMRI representational similarity between trials with more similar orientations of stimulus location in the occipital cortex, such as right V1, right V2, right V3, bilateral POS1, and right lateral occipital 2 (LO2) areas (Bonferroni corrected *p*s < 0.05). We also found the same effect in the oculomotor related region, i.e., the left frontal eye field (FEF), and other regions including the left 31pv, 6ma, 7PC, 8BL, AIP, TE1p, IP2, right 5m, TF, PHT, A4 and parietal area F (PF) (Fig. 6A). Then we tested if any of these brain regions were related to the conflict representation by comparing their encoding strength between incongruent and congruent conditions. Results showed that the right V1, right V2, right POS1, left IP2, left FEF, and right PF encoded stronger orientation effect in the incongruent than the congruent condition, FDR corrected *p*s < .05, one-tailed (Table1, Fig. 6B). We then tested if any of these regions was related to the behavioral performance, and results showed that none of them positively correlated with the behavioral conflict similarity modulation effect, all uncorrected *ps* > .18, one-tailed. Thus all identified regions are consistent with the criterion 3. Like the right 8C area, none of the reported areas directly encoded congruency (see Note S6). Taken together, we found that the visual and oculomotor regions encoded orientations of stimulus location in a continuous manner and that the encoding strength was stronger when the conflict was present.

**Fig. 6.**
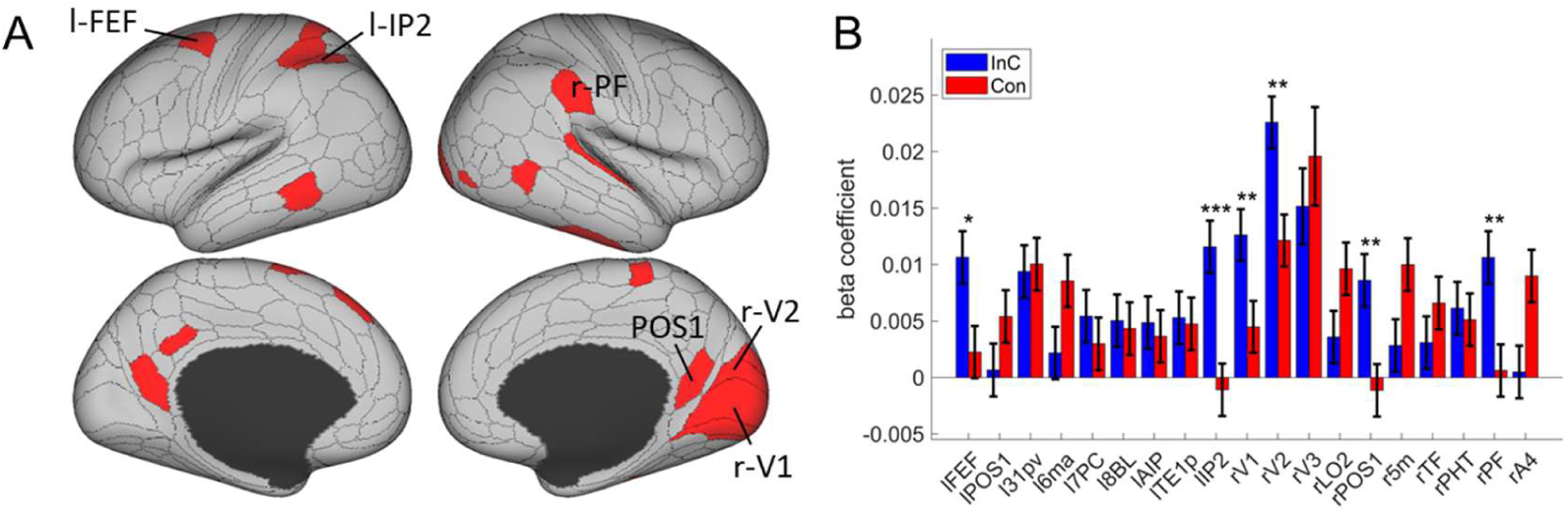
The orientation effect. (A) Brain regions surviving the Bonferroni correction (*p* < 0.05) across the 360 regions (criterion 1). Labeled regions are those meeting the three criterion. (B) Different encoding of orientation in the incongruent with congruent conditions. * FDR corrected *p* < .05, ** FDR corrected *p* < .01; *** FDR corrected *p* < .001.

We hypothesize that the overlapping spatial information of orientation may have facilitated the encoding of conflict types. To explore the relation between conflict type and orientation representations, we conducted representational connectivity (i.e., the similarity between two within-subject RSMs of two regions) (Kriegeskorte et al., 2008) analyses and found that among the orientation effect regions, the right V1 and right V2 showed significant representational connectivity with the right 8C compared to the controlled regions (including those encoding orientation effect but not showing larger encoding strength in incongruent than congruent conditions, as well as eight other regions encoding none of our defined effects in the main RSA, see Methods). Compared with the largest connectivity strength in the controlled regions (i.e., the left 6ma, *β =* 0.7991 ± 0.0299), we found higher connectivity in the right V1, *β =* 0.8633 ± 0.0325, and right V2, *β =* 0.8772 ± 0.0335 (Fig. S6).

## Discussion

Understanding how different types of conflicts are resolved is essential to answer how cognitive control achieves adaptive behavior. However, the dichotomy between domain-general and/or domain-specific processes presents a dilemma (Braem et al., 2014; Egner, 2008). Reconciliation of the two views also suffers from the inability to fully address how different conflicts can be resolved by a limited set of cognitive control processes. In this study, we hypothesized that this issue can be addressed if conflicts are organized as a cognitive space. Leveraging the well-known dissociation between the spatial Stroop and Simon conflicts (Li et al., 2014; Liu et al., 2010; Wang et al., 2014), we designed five conflict types that are systematically different from each other. The cognitive space hypothesis predicted that the representational proximity/distance between two conflict types scales with their similarities/dissimilarities, which was tested at both behavioral and neural levels. Behaviorally, we found that the CSEs were linearly modulated by conflict similarity between consecutive trials, replicating and extending our previous study (Yang et al., 2021). BOLD activity patterns in the right dlPFC further showed that the representational similarity between conflict types was modulated by their conflict similarity, and that strength of the modulation was positively associated with the modulation of conflict similarity on the behavioral CSE. We also observed that activity in two brain regions (right IPS and right dlPFC) was parametrically modulated by the conflict type difference, though they did not directly explain the behavioral results. Additionally, we found that the visual regions encoded the spatial orientation of the stimulus location, which might provide the essential concrete information to determine the conflict type. Together, these results suggest that conflicts may be organized in a cognitive space that enables a limited set of cognitive control processes to resolve a wide variety of distinct types of conflicts.

Conventionally, the domain-general view of control suggests a common representation for different types of conflicts (Fig. 7, left), while the domain-specific view suggests dissociated representations for different types (Fig. 7, right). Previous research on this topic often adopts a binary manipulation of conflicts (Braem et al., 2014) (i.e., each domain only has one conflict type) and gathered evidence for the domain-general/specific view with presence/absence of CSE, respectively. Here, we parametrically manipulated the similarity of conflict types and found the CSE systematically vary with conflict similarity level, demonstrating that cognitive control is neither purely domain-general nor purely domain-specific, but can be reconciled as a cognitive space (Bellmund et al., 2018)(Fig. 7, middle). The model comparison analysis also showed that the Cognitive-Space model explained the representation in right DLPFC better than the Domain-General or Domain-Specific models. Specifically, the cognitive space provides a solution to use a single cognitive space organization to encode different types of conflicts that are close (domain-general) or distant (domain-specific) to each other. It also shows the potential for how various conflict types can be coded using limited resources (i.e., as different points in a low-dimensional cognitive space), as suggested by its out-of-sample predictability. Moreover, geometry can also emerge in the cognitive space (Fu et al., 2022), which will allow for decomposition of a conflict type (e.g., how much conflict in each of the dimensions in the cognitive space) so that it can be mapped into the limited set of cognitive control processes. Such geometry enables fast learning of cognitive control settings from similar conflict types by providing a measure of similarity (e.g., as distance in space).

**Fig. 7.**
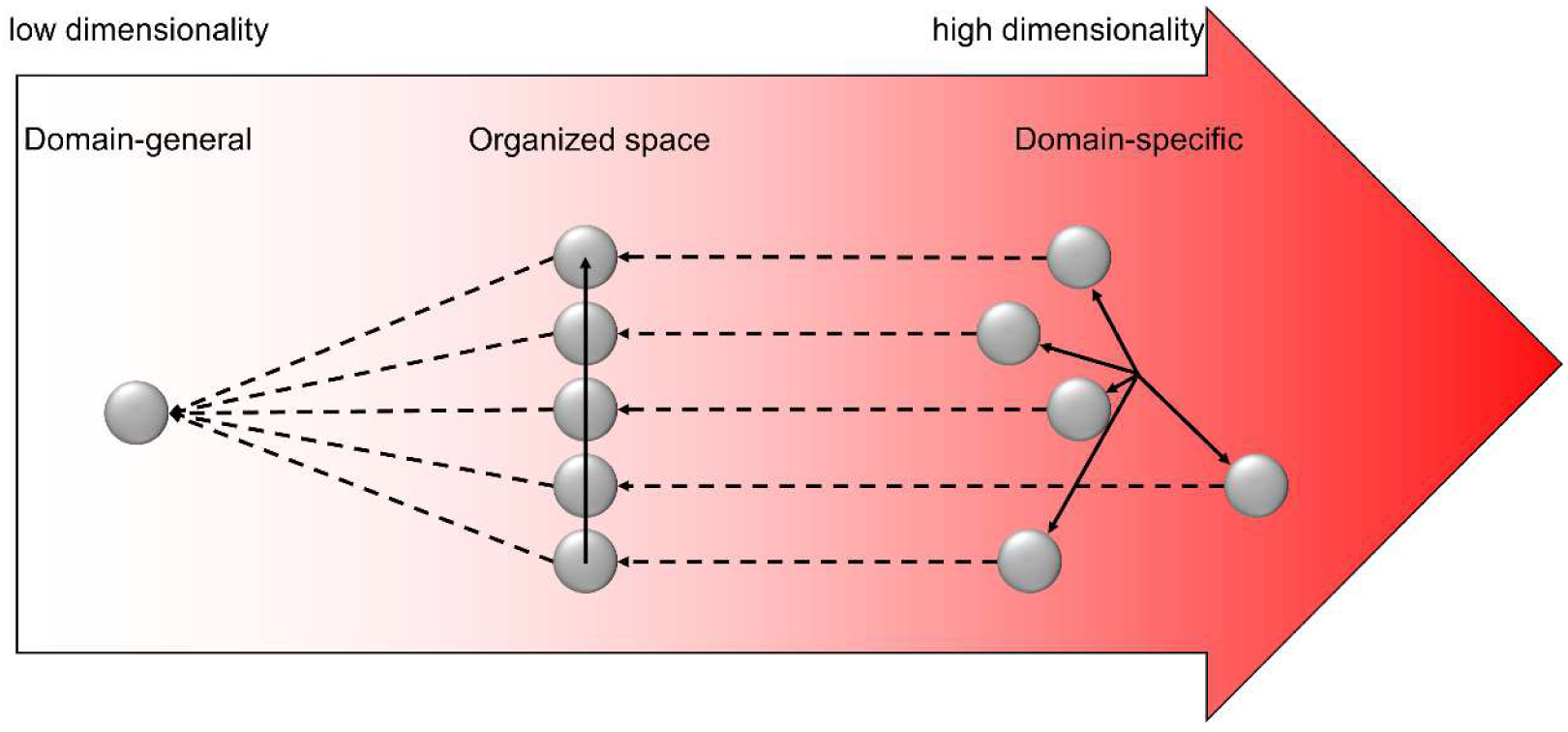
Illustration of the hypothesized dimensionalities of different representations. The shade of the red color indicates the degree of dimensionality (i.e., how many dimensions are needed to represent different states). The dimensionality of domain-general representation is extremely low, with all representations compressed to one dot. The dimensionality of domain-specific representation is extremely high, with each control state encoded in a unique and orthogonal dimension. The dimensionality of the organized representation is modest, enabling distant states to be separated but also allowing close states to share representations. The solid arrows show the axes of different dimensions. The dashed arrows indicate how the representational dimensionality can be reduced by projecting the independent dimensions to a common dimension.

If the dimensionality of the cognitive space of conflict is extremely high, the cognitive space solution would suffer the same criticism as the domain-specificity theory. We argue that the dimensionality is manageable for the human brain, as task information unrelated to differentiating conflicts can be removed. For example, the Simon conflict can be represented in a space consisting of spatial location, stimulus information and responses. Thus, the dimensionality of the cognitive space of conflicts should not exceed the number of represented features. The dimensionality can be further reduced, as humans selectively represent a small number of features when learning task representations (e.g., spatial information is reduced to the horizontal dimension from the 3D space we live in)(Niv, 2019). Consistently, we observed a low dimensional (1.19D) space representing the five conflict types. This is expected since the only manipulated variable is the angular distance between conflict types. The reduced dimensionality does not only require less effort to represent the conflict, but also facilitates generalization of cognitive control settings among different conflict types (Badre et al., 2021).

Although our finding of cognitive space in the right dlPFC differs from other cognitive space studies (Constantinescu et al., 2016; Park et al., 2020; Schuck et al., 2016) that highlighted the orbitofrontal and hippocampus regions, it is consistent with the cognitive control literature. The prefrontal cortex has long been believed to be a key region of cognitive control representation (Mansouri et al., 2007; Miller & Cohen, 2001; Milner, 1963) and is widely engaged in multiple task demands (Cole et al., 2013; Duncan, 2010). However, it is not until recently that the multivariate representation in this region has been examined. For instance, Vaidya et al.(2021) reported that frontal regions presented latent states that are organized hierarchically. Freund et al.(2021) showed that dlPFC encoded the target and congruency in a typical color-word Stroop task. Taken together, we suggest that the right dlPFC might flexibly encode a variety of cognitive spaces to meet the dynamic task demands. In addition, we found no such representation in the left dlPFC (Note S8), indicating a possible lateralization. Previous studies showed that the left dlPFC was related to the expectancy-related attentional set up-regulation, while the right dlPFC was related to the online adjustment of control (Friehs et al., 2020; Vanderhasselt et al., 2009), which is consistent with our findings. Moreover, the right PFC also represents a composition of single rules (Reverberi et al., 2012), which may explain how the spatial Stroop and Simon types can be jointly encoded in a single space.

Previous researchers have proposed an “expected value of control (EVC)” theory, which posits that the brain can evaluate the cost and benefit associated with executing control for a demanding task, such as the conflict task, and specify the optimal control strength (Shenhav et al., 2013). For instance, Grahek et al. (2022) found that more frequently switching goals when doing a Stroop task were achieved by adjusting smaller control intensity. Our work complements the EVC theory by further investigating the neural representation of different conflict conditions and how these representations can be evaluated to facilitate conflict resolution. We found that different conflict conditions could be efficiently represented in a cognitive space encoded by the right dlPFC, and participants with stronger cognitive space representation have also adjusted their conflict control to a greater extent based on the conflict similarity (Fig 4C). The finding suggests that the cognitive space organization of conflicts guides cognitive control to adjust behavior. Previous studies have shown that participants may adopt different strategies to represent a task, with the model-based strategies benefitting goal-related behaviors more than the model-free strategies (Rmus et al., 2022). Similarly, we propose that cognitive space could serve as a mental model to assist fast learning and efficient organization of cognitive control settings. Specifically, the cognitive space representation may provide a principle for how our brain evaluates the expected cost of switching and the benefit of generalization between states and selects the path with the best cost-benefit tradeoff (Abrahamse et al., 2016; Shenhav et al., 2013). The proximity between two states in cognitive space could reflect both the expected cognitive demand required to transition and the useful mechanisms to adapt from. The closer the two conditions are in cognitive space, the lower the expected switching cost and the higher the generalizability when transitioning between them. With the organization of a cognitive space, a new conflict can be quickly assigned a location in the cognitive space, which will facilitate the development of cognitive control settings for this conflict by interpolating nearby conflicts and/or projecting the location to axes representing different cognitive control processes, thus leading to a stronger CSE when following a more similar conflict condition. On the other hand, without a cognitive space, there would be no measure of similarity between conflicts on different trials, hence limiting the ability of fast learning of cognitive control setting from similar trials.

The cognitive space in the right dlPFC appears to be an abstraction of concrete information from the visual regions. We found that the right V1 and V2 encoded the spatial orientation of the target location (Fig. 6) and showed strong representational connectivity with the right dlPFC (Fig. S6), suggesting that there might be information exchange between these regions. We speculate that the representation of spatial orientation may have provided the essential perceptual information to determine the conflict type (Fig. 1) and thus served as the critical input for the cognitive space. The conflict type representation further incorporates the stimulus-response mapping rules to the spatial orientation representation, so that vertically symmetric orientations can be recognized as the same conflict type (Fig. 7). In other words, the representation of conflict type involves the compression of perceptual information (Flesch et al., 2022), which is consistent with the idea of a low-dimensional representation of cognitive control (Badre et al., 2021; MacDowell et al., 2022). The compression and abstraction processes might be why the frontoparietal regions are the top of hierarchy of information processing (Gilbert & Li, 2013) and why the frontoparietal regions are widely engaged in multiple task demands (Duncan, 2013).

Although the spatial orientation information in our design could be helpful to the construction of cognitive space, the cognitive space itself was independent of the stimulus-level representation of the task. We found the conflict similarity modulation on CSE did not change with more training (see Note S3), indicating that the cognitive space did not depend on strategies that could be learned through training. Instead, the cognitive space should be determined by the intrinsic similarity structure of the task design. For example, a previous study (Freitas & Clark, 2015) has found that the CSE across different versions of spatial Stroop and flanker tasks was stronger than that across either of the two conflicts and Simon. In their designs, the stimulus similarity was controlled at the same level, so the difference in CSE was only attributable to the similar dimensional overlap between Stroop and flanker tasks, in contrast to the Simon task. Furthermore, recent studies showed that the cognitive space generally exists to represent structured latent states (e.g., Vaidya & Badre, 2022), mental strategy cost (Grahek et al., 2022), and social hierarchies (Park et al., 2020). Therefore, cognitive space is likely a universal strategy that can be applied to different scenarios.

With conventional univariate analyses, we observed that the overall congruency effect was located at the medial frontal region (i.e., pre-SMA), which is consistent with previous studies (Botvinick et al., 2004; Fu et al., 2022). Beyond that, we also found regions that can be parametrically modulated by conflict type difference, including right IPS, right dlPFC (modulated by Simon difference). The right lateralization of these regions is consistent with a previous finding (Li et al., 2017). The parametric encoding of conflict also mirrors prior research showing the parametric encoding of task demands (Dagher et al., 1999; Ritz & Shenhav, 2023). The scaling of brain activities based on conflict difference is potentially important to the representational organization of different types of conflicts. However, we didn’t observe their brain-behavioral relevance. One possible reason is that the conflict (dis)similarity is a combination of (dis)similarity of spatial Stroop and Simon conflicts, but each univariate region only reflects difference along a single conflict domain. Also likely, the representational geometry is more of a multivariate problem than what univariate activities can capture (Freund, Etzel, et al., 2021). Future studies may adopt approaches such as repetition suppression induced fMRI adaptation (Badre et al., 2021) to test the role of univariate activities in task representations.

Recently an interesting debate has arisen concerning whether cognitive control should be considered as a process or a representation (Freund, Etzel, et al., 2021). Traditionally, cognitive control has been predominantly viewed as a process. However, the study of its representation has gained more and more attention. While it may not be as straightforward as the visual representation (e.g., creating a mental image from a real image in the visual area), cognitive control can have its own form of representation. An influential theory, Marr’s (1982) three-level model proposed that representation serves as the algorithm of the process to achieve a goal based on the input. In other words, representation can encompass a dynamic process rather than being limited to static stimuli. Building on this perspective, we posit that the representation of cognitive control consists of an array of dynamic representations embedded within the overall process. A similar idea has been proposed that the representation of task profiles can be progressively constructed with time in the brain (Kikumoto & Mayr, 2020). Moreover, we anticipate that the representation of cognitive space is most prominently involved at two critical stages to guide the transference of behavioral CSE. The first stage involves the evaluation of control demands, where the representational distance/similarity between previous and current trials influences the adjustment of cognitive control. The second stage pertains to control execution, where the switch from one control state to another follows a path within the cognitive space. However, we were unable to fully distinguish between these two stages due to the low temporal resolution of fMRI signals in our study. Future research seeking to delve deeper into this question may benefit from methodologies with higher temporal resolutions, such as EEG and MEG.

Several interesting questions remains to be answered. For example, is the dimension of the unified space across conflict-inducing tasks solely determined by the number of conflict sources? Is this unified space adaptively adjusted within the same brain region? Can we effectively map any sources of conflict with completely different stimuli into a single space? Does the cognitive space vary from population to population, such as between the normal people and patients?

Methodological implications. Previous studies with mixed conflicts have applied mainly categorical manipulations of conflict types, such as the multi-source interference task (Fu et al., 2022) and color Stroop-Simon task (Liu et al., 2010). The categorical manipulations make it difficult to quantify conceptual similarity between conflict types and hence limit the ability to test whether neural representations of conflict capture conceptual similarity. To the best of our knowledge, no previous studies have manipulated the conflict types parametrically. This gap highlights a broader challenge within cognitive science: effectively manipulating and measuring similarity levels for conflicts, as well as other high-level cognitive processes, which are inherently abstract. The use of an experimental paradigm that permits parametric manipulation of conflict similarity provides a way to systematically investigate the organization of cognitive control, as well as its influence on adaptive behaviors. Moreover, the cross-subject RSA provides high sensitivity to the variables of interest and the ability to separate confounding factors. For instance, in addition to dissociating conflict type from orientation, we dissociated target from response, and spatial Stroop distractor from Simon distractor. We further showed cognitive control can both enhance the target representation and suppress the distractor representation (Note S10, Fig. S7), which is in line with previous studies (Polk et al., 2008; Ritz & Shenhav, 2022).

### Limitations

A few limitations of this study need to be noted. To parametrically manipulate the conflict similarity levels, we adopted the spatial Stroop-Simon paradigm that enables parametrical combinations of spatial Stroop and Simon conflicts. However, since this paradigm is a two-alternative forced choice design, the behavioral CSE is not a pure measure of adjusted control but could be partly confounded by bottom-up factors such as feature integration (Hommel et al., 2004). Future studies may replicate our findings with a multiple-choice design (including more varied stimulus sets, locations and responses) with confound-free trial sequences (Braem et al., 2019). Another limitation is that in our design, the spatial Stroop and Simon effects are highly anticorrelated. This constraint may make the five conflict types represented in a unidimensional space (e.g., a circle) embedded in a 2D space. This limitation also means we cannot conclusively rule out the possibility of a real unidimensional space driven solely by spatial Stroop or Simon conflicts. However, this appears unlikely, as it would imply that our manipulation of conflict types merely represented varying levels of a single conflict, akin to manipulating task difficulty when everything else being equal. If task difficulty were the primary variable, we would expect to see greater representational similarity between task conditions of similar difficulty, such as the Stroop and Simon conditions, which demonstrates comparable congruency effects (see Fig. S1). Contrary to this, our findings reveal that the Stroop-only and Simon-only conditions exhibit the lowest representational similarity (Fig. S4). Furthermore, Fu et al. (2022) has shown that the representation of mixtures of Simon and Flanker conflicts was compositional, rather than reflecting single dimension, which also applies to our cases. Future studies may test the 2D cognitive space with fully independent conditions. A possible improvement to our current design would be to include left, right, up, and down arrows represented in a grid formation across four spatially separate quadrants, with each arrow mapped to its own response button. Additionally, our study is not a comprehensive test of the cognitive space hypothesis but aimed primarily to provide original evidence for the geometry of cognitive space in representing conflict information in cognitive control. Future research should examine other aspects of the cognitive space such as its dimensionality, its applicability to other conflict tasks such as Eriksen Flanker task, and its relevance to other cognitive abilities, such as cognitive flexibility and learning.

In sum, we showed that the cognitive control can be parametrically encoded in the right dlPFC and guides cognitive control to adjust goal-directed behavior. This finding suggests that different cognitive control states may be encoded in an abstract cognitive space, which reconciles the long-standing debate between the domain-general and domain-specific views of cognitive control and provides a parsimonious and more broadly applicable framework for understanding how our brains efficiently and flexibly represents multiple task settings.

## Materials and Methods

### Subjects

In Experiment 1, we enrolled thirty-three college students (ages 19-28, average 21.5 ± 2.3 years; 19 males). In Experiment 2, thirty-six college students were recruited, one of which was excluded due to not following task instructions. The final sample of Experiment 2 consisted of thirty-five participants (ages 19-29, average 22.3 ± 2.5 years; 17 males). The sample sizes were determined based on our previous study (Yang et al., 2021). All participants reported no history of psychiatric or neurological disorders and were right-handed, with normal or corrected-to-normal vision. The experiments were approved by the Institutional Review Board of the Institute of Psychology, Chinese Academy of Science. Informed consent was obtained from all subjects.

### Method Details

#### Experiment 1

##### Experimental Design

We adopted a modified spatial Stroop-Simon task (Yang et al., 2021) (Fig. 1). The task was programmed with E-prime 2.0 (Psychological Software Tools, Inc.). The stimulus was an upward or downward black arrow (visual angle of ∼ 1°), displayed on a 17-inch LCD monitor with a viewing distance of ∼60 cm. The arrow appeared inside a grey square at one of ten locations with the same distance from the center of the screen, including two horizontal (left and right), two vertical (top and bottom), and six corner (orientations of 22.5°, 45° and 67.5°) locations. The distance from the arrow to the screen center was approximately 3°. To dissociate orientation of stimulus locations and conflict types (see below), participants were randomly assigned to two sets of stimulus locations (one included top-right and bottom-left quadrants, and the other included top-left and bottom-right quadrants).

Each trial started with a fixation cross displayed in the center for 100−300 ms, followed by the arrow for 600 ms and another fixation cross for 1100−1300 ms (the total trial length was fixed at 2000 ms). Participants were instructed to respond to the pointing direction of the arrow by pressing a left or right button and to ignore its location. The mapping between the arrow orientations and the response buttons was counterbalanced across participants. The task design introduced two possible sources of conflicts: on one hand, the direction of the arrow is either congruent or incongruent with the vertical location of the arrow, thus introducing a spatial Stroop conflict (Lu & Proctor, 1995; MacLeod, 1991), which contains the dimensional overlap between task-relevant stimulus and task-irrelevant stimulus (Kornblum et al., 1990); on the other hand, the response (left or right button) is either congruent or incongruent with the horizontal location of the arrow, thus introducing a Simon conflict (Lu & Proctor, 1995; Simon & Small, 1969), which contains the dimensional overlap between task-irrelevant stimulus and response (Kornblum et al., 1990). Therefore, the five polar orientations of the stimulus location (from 0 to 90°) defined five unique combinations of spatial Stroop and Simon conflicts, with more similar orientations having more similar composition of conflicts. More generally, the spatial orientation of the arrow location relative to the center of the screen forms a cognitive space of different blending of spatial Stroop and Simon conflicts.

The formal task consisted of 30 runs of 101 trials each, divided into three sessions of ten runs each. The participants completed one session each time and all three sessions within one week. Before each session, the participants performed training blocks of 20 trials repeatedly until the accuracy reached 90% in the most recent block. The trial sequences of the formal task were pseudo-randomly generated to ensure that each of the task conditions and their transitions occurred with equal number of trials.

#### Experiment 2

##### Experimental Design

The apparatus, stimuli and procedure were identical to Experiment 1 except for the changes below. The stimuli were back projected onto a screen (with viewing angle being ∼3.9° between the arrow and the center of the screen) behind the subject and viewed via a surface mirror mounted onto the head coil. Due to the time constraints of fMRI scanning, the trial numbers decreased to a total of 340, divided into two runs with 170 trials each. To obtain a better hemodynamic model fitting, we generated two pseudo-random sequences optimized with a genetic algorithm (Wager & Nichols, 2003) conducted by the NeuroDesign package (Durnez et al., 2018) (see Note S3 for more detail). In addition, we added 6 seconds of fixation before each run to allow the stabilization of the hemodynamic signal, and 20 seconds after each run to allow the signal to drop to the baseline.

Before scanning, participants performed two practice sessions. The first one contained 10 trials of center-displayed arrow and the second one contained 32 trials using the same design as the main task. They repeated both sessions until their performance accuracy for each session reached 90%, after which the scanning began.

### fMRI Image acquisition and preprocessing

Functional imaging was performed on a 3T GE scanner (Discovery MR750) using echo-planar imaging (EPI) sensitive to BOLD contrast [in-plane resolution of 3.5 × 3.5 mm^2^, 64 × 64 matrix, 37 slices with a thickness of 3.5 mm and no interslice skip, repetition time (TR) of 2000 ms, echo-time (TE) of 30 ms, and a flip angle of 90°]. In addition, a sagittal T1-weighted anatomical image was acquired as a structural reference scan, with a total of 256 slices at a thickness of 1.0 mm with no gap and an in-plane resolution of 1.0 × 1.0 mm^2^.

Before preprocessing, the first three volumes of the functional images were removed due to the instability of the signal at the beginning of the scan. The anatomical and functional data were preprocessed with the fMRIprep 20.2.0 (Esteban et al., 2019) (RRID:SCR_016216), which is based on Nipype 1.5.1 (Gorgolewski et al., 2011) (RRID:SCR_002502). Specifically, BOLD runs were slice-time corrected using 3dTshift from AFNI 20160207 (Jenkinson et al., 2002) (RRID:SCR_005927). The BOLD time-series were resampled to the MNI152NLin2009cAsym space without smoothing. For a more detailed description of preprocessing, see Note S4. After preprocessing, we resampled the functional data to a spatial resolution of 3 × 3 × 3 mm^3^. All analyses were conducted in volumetric space, and surface maps are produced with Connectome Workbench (https://www.humanconnectome.org/software/connectome-workbench) for display purpose only.

### Quantification and Statistical Analysis

#### Behavioral analysis

##### Experiment 1

RT and ER were the two dependent variables analyzed. As for RTs, we excluded the first trial of each block (0.9%, for CSE analysis only), error trials (3.8%), trials with RTs beyond three *SD*s or shorter than 200 ms (1.3%) and post-error trials (3.4%). For the ER analysis, the first trial of each block and trials after an error were excluded. To exclude the possible influence of response repetition, we centered the RT and ER data within the response repetition and response alternation conditions separately by replacing condition-specific mean with the global mean for each subject.

To examine the modulation of conflict similarity on the CSE, we organized trials based on a 5 (previous trial conflict type) × 5 (current trial conflict type) × 2 (previous trial congruency) × 2 (current trial congruency) factorial design. As conflict similarity is commutive between conflict types, we expected the previous by current trial conflict type factorial design to be a symmetrical (e.g., a conflict 1-conflict 2 sequence in theory has the same conflict similarity modulation effect as a conflict 2-conflict 1 sequence), resulting a total of 15 conditions left for the first two factors of the design (i.e., previous × current trial conflict type). For each previous × current trial conflict type condition, the conflict similarity between the two trials can be quantified as the cosine of their angular difference. In the current design, there were five possible angular difference levels (0, 22.5°, 42.5°, 67.5° and 90°, see Fig. 1C). We further coded the previous by current trial congruency conditions (hereafter abbreviated as CSE conditions) as CC, CI, IC and II, with the first and second letter encoding the congruency (C) or incongruency (I) on the previous and current trial, respectively. As the CSE is operationalized as the interaction between previous and current trial congruency, it can be rewritten as a contrast of (CI – CC) – (II – IC). In other words, the load of CSE on CI, CC, II and IC conditions is 1, –1, –1 and 1, respectively. To estimate the modulation of conflict similarity on the CSE, we built a regressor by calculating the Kronecker product of the conflict similarity scores of the 15 previous × current trial conflict similarity conditions and the CSE loadings of previous × current trial congruency conditions. This regressor was regressed against RT and ER data separately, which were normalized across participants and CSE conditions. The regression was performed using a linear mixed-effect model, with the intercept and the slope of the regressor for the modulation of conflict similarity on the CSE as random effects (across both participants and the four CSE conditions). As a control analysis, we built a similar two-stage model (Yang et al., 2021). In the first stage, the CSE [i.e., (CI – CC) – (II – IC)] for each of the previous × current trial conflict similarity condition was computed. In the second stage, CSE was used as the dependent variable and was predicted using conflict similarity across the 15 previous × current trial conflict type conditions. The regression was also performed using a linear mixed effect model with the intercept and the slope of the regressor for the modulation of conflict similarity on the CSE as random effects (across participants). *Experiment 2.* Behavioral data was analyzed using the same linear mixed effect model as Experiment 1, with all the CC, CI, IC and II trials as the dependent variable. In addition, to test if fMRI activity patterns may explain the behavioral representations differently in congruent and incongruent conditions, we conducted the same analysis to measure behavioral modulation of conflict similarity on the CSE using congruent (CC and IC) and incongruent (CI and II) trials separately.

### Estimation of fMRI activity with univariate general linear model (GLM)

To estimate voxel-wise fMRI activity for each of the experimental conditions, the preprocessed fMRI data of each run were analyzed with the GLM. We conducted three GLMs for different purposes. GLM1 aimed to validate the design of our study by replicating the engagement of frontoparietal activities in conflict processing documented in previous studies (Jiang & Egner, 2014; Li et al., 2017), and to explore the cognitive space related regions that were parametrically modulated by the conflict type. Preprocessed functional images were smoothed using a 6-mm FWHM Gaussian kernel. We included incongruent and congruent conditions as main regressors and appended a parametric modulator for each condition. The modulation parameters for Stroop, St_H_Sm_L_, St_M_Sm_M_, St_L_Sm_H_, and Simon trials were −2, −1, 0, 1 and 2, respectively. In addition, we also added event-related nuisance regressors, including error/missed trials, outlier trials (slower than three SDs of the mean or faster than 200 ms) and trials within two TRs of significant head motion (i.e., outlier TRs, defined as standard DVARS > 1.5 or FD > 0.9 mm from previous TR)(Jiang et al., 2020). On average there were 1.2 outlier TRs for each run. These regressors were convolved with a canonical hemodynamic response function (HRF) in SPM 12 (http://www.fil.ion.ucl.ac.uk/spm). We further added volume-level nuisance regressors, including the six head motion parameters, the global signal, the white matter signal, the cerebrospinal fluid signal, and outlier TRs. Low-frequency signal drifts were filtered using a cutoff period of 128 s. The two runs were regarded as different sessions and incorporated into a single GLM to get more power. This yielded two beta maps (i.e., a main effect map and a parametric modulation map) for the incongruent and congruent conditions, respectively and for each subject. At the group level, paired t-tests were conducted between incongruent and congruent conditions, one for the main effect and the other for the parametric modulation effect. Since the spatial Stroop and Simon conflicts change in the opposite direction to each other, a positive modulation effect would reflect a higher brain activation when there is more Simon conflict, and a negative modulation effect would reflect a higher brain activation for more spatial Stroop conflict. To avoid confusion, we converted the modulation effect of spatial Stroop to positive by using a contrast of [– (I_pm – C_pm)] throughout the results presentation. Results were corrected with the probabilistic threshold-free cluster enhancement (pTFCE) and then thresholded by 3dClustSim function in AFNI (Cox & Hyde, 1997) with voxel-wise *p* < .001 and cluster-wize *p* < .05, both 1-tailed. To visualize the parametric modulation effects, we conducted a similar GLM (GLM2), except we used incongruent and congruent conditions from each conflict type as separate regressors with no parametric modulation. Then we extracted beta coefficients for each regressor and each participant with regions observed in GLM1 as regions of interest, and finally got the incongruent−congruent contrasts for each conflict type at the individual level. We reported the results in Fig. 3, Table S1, and Fig. S3. Visualization of the uni-voxel results was made by the MRIcron (https://www.mccauslandcenter.sc.edu/mricro/mricron/).

The GLM3 aimed to prepare for the representational similarity analysis (see below). There were several differences compared to GLM1. The unsmoothed functional images after preprocessing were used. This model included 20 event-related regressors, one for each of the 5 (conflict type) × 2 (congruency condition) × 2 (arrow direction) conditions. The event-related nuisance regressors were similar to GLM1, but with additional regressors of response repetition and post-error trials to account for the nuisance inter-trial effects. To fully expand the variance, we conducted one GLM analysis for each run. After this procedure, a voxel-wise fMRI activation map was obtained per condition, run and subject.

### Representational similarity analysis (RSA)

To measure the neural representation of conflict similarity, we adopted the RSA. RSAs were conducted on each of the 360 cortical regions of a volumetric version of the MMP cortical atlas (Glasser et al., 2016). To de-correlate the factors of conflict type and orientation of stimulus location, we leveraged the between-subject manipulation of stimulus locations and conducted RSA in a cross-subject fashion (Fig. 4). Previous studies (e.g., J. Chen et al., 2017) have demonstrated that consistent multi-voxel activation patterns exist across individuals, and successful applications of cross-subject RSA (see review by Freund, Etzel, et al., 2021) and cross-subject decoding approaches (Jiang et al., 2016; Tusche et al., 2016) have also been reported. The beta estimates from GLM3 were noise-normalized by dividing the original beta coefficients by the square root of the covariance matrix of the error terms (Nili et al., 2014). For each cortical region, we calculated the Pearson’s correlations between fMRI activity patterns for each run and each subject, yielding a 1400 (20 conditions × 2 runs × 35 participants) × 1400 RSM. The correlations were calculated in a cross-voxel manner using the fMRI activation maps obtained from GLM3 described in the previous section. We excluded within-subject cells from the RSM (thus also excluding the within-run similarity as suggested by Walther et al., (2016)), and the remaining cells were converted into a vector, which was then z-transformed and submitted to a linear mixed effect model as the dependent variable. The linear mixed effect model also included regressors of conflict similarity and orientation similarity. Importantly, conflict similarity was based on how Simon and spatial Stroop conflicts are combined and hence was calculated by first rotating all subject’s stimulus location to the top-right and bottom-left quadrants, whereas orientation was calculated using original stimulus locations. As a result, the regressors representing conflict similarity and orientation similarity were de-correlated (Fig. 4A). Similarity between two conditions was measured as the cosine value of the angular difference. Other regressors included a target similarity regressor (i.e., whether the arrow directions were identical), a response similarity regressor (i.e., whether the correct responses were identical); a spatial Stroop distractor regressor (i.e., vertical distance between two stimulus locations); a Simon distractor regressor (i.e., horizontal distance between two stimulus locations). Additionally, we also included a regressor denoting the similarity of Group (i.e., whether two conditions are within the same subject group, according to the stimulus-response mapping). We also added two regressors including ROI-mean fMRI activations for each condition of the pair to remove the possible uni-voxel influence on the RSM. A last term was the intercept. To control the artefact due to dependence of the correlation pairs sharing the same subject, we included crossed random effects (i.e., row-wise and column-wise random effects) for the intercept, conflict similarity, orientation and the group factors (G. Chen et al., 2017). Individual effects for each regressor were also extracted from the model for brain-behavioral correlation analyses. In brain-behavioral analyses, only the RT was used as behavioral measure to be consistent with the fMRI results, where the error trials were regressed out.

The statistical significance of these beta estimates was based on the outputs of the mixed-effect model estimated with the “fitlme” function in Matlab 2022a. We adjusted the t and p values with the degrees of freedom calculated through the Satterthwaite approximation method (Satterthwaite, 1946). Of note, this approach was applied to all the mixed-effect model analyses in this study. Multiple comparison correction was applied with the Bonferroni approach across all cortical regions at the *p* < 0.05 level. To test if the representation strengths are different between congruent and incongruent conditions, we also conducted the RSA using only congruent (RDM_C) and incongruent (RDM_I) trials separately. The contrast analysis was achieved by an additional model with both RDM_C and RDM_I included, adding the congruency and the interaction between conflict type (and orientation) and congruency as both fixed and random factors. The difference between incongruent and congruent representations was indicated by a significant interaction effect. To visualize the difference, we plotted the effect-related patterns (the predictor multiplied by the slope, plus the residual) as a function of the similarity levels (Fig. 5D), and a summary RSM for incongruent and congruent conditions, respectively (Fig. S4).

### Model comparison and representational dimensionality

To estimate if the right 8C specifically encodes the cognitive space, rather than the domain-general or domain-specific structures, we conducted two more RSAs. We replaced the cognitive space-based conflict similarity matrix in the RSA we reported above (hereafter referred to as the Cognitive-Space model) with one of the alternative model matrices, with all other regressors equal. The domain-general model treats each conflict type as equivalent, so each two conflict types only differ in the magnitude of their conflict. Therefore, we defined the domain-general matrix as the absolute difference in their congruency effects indexed by the group-averaged RT in Experiment 2. Then the z-scored model vector was sign-flipped to reflect similarity instead of distance. The domain-specific model treats each conflict type differently, so we used a diagonal matrix, with within-conflict type similarities being 1 and all cross-conflict type similarities being 0.

To better capture the dimensionality of the representational space, we estimated its dimensionality using the participation ratio (Ito & Murray, 2023). Since we excluded the within-subject cells from the whole RSM, the whole RSM is an incomplete matrix and could not be used. To resolve this issue, we averaged the cells corresponding to each pair of conflict types to obtain an averaged 5×5 RSM matrix, similar to the matrix shown in Fig. 1C. We then estimated the participation ratio using the formula:

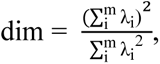

where λ_i_ is the eigenvalue of the RSM and m is the number of eigenvalues.

### Representational connectivity analysis

To explore the possible relevance between the conflict type and the orientation effects, we conducted representational connectivity (Kriegeskorte et al., 2008) between regions showing evidence encoding conflict similarity and orientation similarity. We hypothesized that this relationship should exist at the within-subject level, so we conducted this analysis using within-subject RSMs excluding the diagonal. Specifically, the z-transformed RSM vector of each region were extracted and submitted to a mixed linear model, with the RSM of the conflict type region (i.e., the right 8C) as the dependent variable, and the RSM of one of the orientation regions (e.g., right V2) as the predictor. Intercept and the slope of the regressor were set as random effects at the subject level. The mixed effect model was conducted for each pair of regions, respectively. Considering there might be strong intrinsic correlations across the RSMs induced by the nuisance factors, such as the within-subject similarity, we added two sets of regions as control. First, we selected regions without showing any effects of interest (i.e., uncorrected *ps* > 0.3 for all the conflict type, orientation, congruency, target, response, spatial Stroop distractor and Simon distractor effects). Second, we selected regions of orientation effect meeting the first but not the second criterion, to account for the potential correlation between regions of the two partly orthogonal regressors (Fig. 4A). Regions adjacent to the orientation regions were excluded to avoid the inherent strong similarity they may share. Existence of representational connectivity was defined by a connectivity slope higher than 95% of the standard error above the mean of any control region.

## Acknowledgement

We thank Eliot Hazeltine for valuable comments on a previous version of this manuscript. The work was supported by the National Natural Science Foundation of China and the German Research Foundation (NSFC 62061136001/DFG TRR-169) to X.L. and China Postdoctoral Science Foundation (2019M650884) to G.Y.

