## Supplementary Material_R3 for "Conflicts are parametrically encoded: initial evidence for a cognitive space view to reconcile the debate of domain-general and domain-specific cognitive control"

### Supplementary Notes

***Note S1.*** *Behavioral congruency effects*

To test the congruency effects for the five conflict types, we conducted 5 (conflict type) × 2 (congruency) repeated-measure ANOVAs with RT and ER from both experiments. The results are displayed in Supplementary Fig. 1.

*Experiment 1.*

For the RT, we observed a significant main effect of Congruency, $F(1, 32) = 407.70$, $p < .001$, $\eta_p^2 = .93$, a significant main effect of Conflict Type, $F(4, 128) = 6.32$, $p < .001$, $\eta_p^2 = .16$, and an interaction between Conflict Type and Congruency, $F(4, 128) = 27.86$, $p < .001$, $\eta_p^2 = .47$. Simple effect analyses showed that participants responded more slowly in incongruent conditions than in congruent conditions for all conflict types, $p_{FDR}s < .001$. Additionally, the congruency effect of the Type 2, 3 and 4 were significantly larger than that of the Type 1, and the congruency effect of the Type 2 and 3 were significantly larger than that of the Type 5, $p_{FDR}$s < .05.

Similar results were found with the ER. We observed a significant main effect of Congruency, $F(1, 32) = 56.83$, $p < .001$, $\eta_p^2 = .64$, a significant main effect of Conflict Type, $F(4, 128) = 6.29$, $p < .001$, $\eta_p^2 = .16$, and an interaction between Conflict Type and Congruency, $F(4, 128) = 13.23$, $p < .001$, $\eta_p^2 = .29$. Simple effect analyses showed that participants were more error-prone in incongruent conditions than in congruent conditions for all conflict types, $p_{FDR}s < .001$. The congruency effect of the Type 2, 3 and 4 were significantly larger than that of the Type 1, and the congruency effect of the Type 3 and 4 were significantly larger than that of the Type 5, $p_{FDR}$s < .05.

*Experiment 2.*

For the RT, we observed a significant main effect of Congruency, $F(1, 34) = 149.71$, $p < .001$, $\eta_p^2 = .81$, a significant main effect of Conflict Type, $F(4, 136) = 10.11$, $p < .001$, $\eta_p^2 = .23$, and an interaction between Conflict Type and Congruency, $F(4, 136) = 7.63$, $p < .001$, $\eta_p^2 = .18$. Simple effect analyses showed that participants responded more slowly in incongruent conditions than in congruent conditions for all conflict types, $p_{FDR}$s < .001. The congruency effect of the Type 4 condition was larger than that of Type 1, and Type 3 and Type 4 were significantly larger than that of the Type 5, $p_{FDR}$s < .05.

For the ER, we only observed a significant main effect of Congruency, $F(1, 34) = 29.80$, $p < .001$, $\eta_p^2 = .47$. All the types showed a larger error rate in incongruent than congruent conditions ($p_{FDR}$s < .001), except that the Type 1 only showed a marginal significance ($p_{FDR} = .062$).

In sum, we observed strong behavioral congruency effects in both experiments. The findings indicate that these conflict conditions indeed engaged cognitive control (Freund, Etzel, et al., 2021).

***Note S2.*** *Modulation of conflict similarity on behavioral CSEs cannot be explained by the physical proximity*

In our design, the conflict similarity might be confounded by the physical proximity between stimulus (i.e., the arrow) of two consecutive trials. That is, when arrows of the two trials appear at the same quadrant, a higher conflict similarity also indicates a higher physical proximity (Fig. 1A). Although the opposite is true if arrows of the two trials appear at different quadrants, it is possible the behavioral effects can be biased by the within quadrant trials. To examine if the physical distance has confounded the conflict similarity modulation effect, we conducted an additional analysis.

We defined the physical angular difference across two trials as the difference of their polar angles relative to the origin. Therefore, the physical angular difference could vary from 0 to 180°. For each CSE conditions (i.e., CC, CI, IC and II), we grouped the trials based on their physical angular distances, and then averaged trials with the same previous by current conflict type transition but different orders (e.g., $St_HSm_L$−$St_LSm_H$ and $St_LSm_H$−$St_HSm_L$) within each subject. The data were submitted to a mixed-effect model with the conflict similarity, physical proximity (i.e., the opposite of the physical angular difference) as fixed-effect predictors, and subject and CSE condition as random effects. Results showed significant conflict similarity modulation effects in both Experiment 1 (RT: $\beta = 0.09 \pm 0.01$, $t(1902.4) = 13.74$, $p < .001$, $\eta_p^2 = .025$; ER: $\beta = 0.09 \pm 0.01$, $t(249.3) = 7.66$, $p < .001$, $\eta_p^2 = .018$) and Experiment 2 (RT: $\beta = 0.21 \pm 0.02$, $t(61.0) = 4.71$, $p < .001$, $\eta_p^2 = .056$; ER: $\beta = 0.20 \pm 0.03$, $t(65.0) = 4.16$, $p < .001$, $\eta_p^2 = .236$). Thus, the observed modulation of conflict similarity on behavioral CSEs cannot be explained by physical proximity.

***Note S3.*** *Modulation of conflict similarity on behavioral CSEs does not change across time*

We tested if the conflict similarity modulation on the CSE is susceptible to training. We collected the data of Experiment 1 across three sessions, thus it is possible to examine if the conflict similarity modulation effect changes across time. To this end, we added conflict similarity, session and their interaction into a mixed-effect linear model, in which the session was set as a categorical variable. With a post-hoc analysis of variance (ANOVA), we calculated the statistical significance of the interaction term. This approach was applied to both the RT and ER. Results showed no interaction effect in either RT, $F(2,76.4) = 1.025$, $p = .364$, or ER, $F(2,49.4) = 0.789$, $p = .460$. This result suggests that the modulation effect does not change across time.

***Note S4.*** *The fMRI sequence generation approach*

Two sequences of 170 trials each were generated independently with the NeuroDesign package (Durnez et al., 2018). Each sequence was initialized as 10 consecutive sub-blocks of each condition (incongruent and congruent) for each conflict type (Stroop, $St_HSm_L$, $St_MSm_M$, $St_LSm_H$, and Simon). The contrasts of interest were the main effect of congruency (i.e., [1 −1 1 −1 1 −1 1 −1 1 −1]) and the parametric effect (i.e., [−2 −2 −1 −1 0 0 1 1 2 2]). The order was optimized after 5000 cycles of crossover, mutation, immigration, fitness, and natural selection. The final number of trials for different conflict types varied from 64 to 73.

***Note S5.*** *fMRI data preprocessing*

Results included in this manuscript come from preprocessing performed using fMRIPrep 20.2.0 (RRID:SCR_016216)(Esteban et al., 2019), which is based on Nipype 1.5.1 (RRID:SCR_002502)(Gorgolewski et al., 2011).

*Anatomical data preprocessing.* The T1-weighted (T1w) image was corrected for intensity non-uniformity (INU) with N4BiasFieldCorrection (Tustison et al., 2010), distributed with ANTs 2.3.3 (RRID:SCR_004757)(Avants et al., 2008), and used as T1w-reference throughout the workflow. The T1w-reference was then skull-stripped with a Nipype implementation of the antsBrainExtraction.sh workflow (from ANTs), using OASIS30ANTs as target template. Brain tissue segmentation of cerebrospinal fluid (CSF), white-matter (WM) and gray-matter (GM) was performed on the brain-extracted T1w using fast (FSL 5.0.9, RRID:SCR_002823)(Zhang et al., 2001). Volume-based spatial normalization to one standard space (MNI152NLin2009cAsym) was performed through nonlinear registration with antsRegistration (ANTs 2.3.3), using brain-extracted versions of both T1w reference and the T1w template. The following template was selected for spatial normalization: ICBM 152 Nonlinear Asymmetrical template version 2009c [RRID:SCR_008796; TemplateFlow ID: MNI152NLin2009cAsym](Fonov et al., 2009).

*Functional data preprocessing.* For each of the 5 BOLD runs found per subject (across all tasks and sessions), the following preprocessing was performed. First, a reference volume and its skull-stripped version were generated using a custom methodology of fMRIPrep. Susceptibility distortion correction (SDC) was omitted. The BOLD reference was then co-registered to the T1w reference using flirt (FSL 5.0.9)(Jenkinson & Smith, 2001) with the boundary-based registration (Greve & Fischl, 2009) cost-function. Co-registration was configured with nine degrees of freedom to account for distortions remaining in the BOLD reference. Head-motion parameters with respect to the BOLD reference (transformation matrices, and six corresponding rotation and translation parameters) are estimated before any spatiotemporal filtering using mcflirt (FSL 5.0.9)(Jenkinson et al., 2002). BOLD runs were slice-time corrected using 3dTshift from AFNI 20160207

(RRID:SCR_005927)(Jenkinson et al., 2002). The BOLD time-series (including slice-timing correction when applied) were resampled onto their original, native space by applying the transforms to correct for head-motion. These resampled BOLD time-series will be referred to as preprocessed BOLD in original space, or just preprocessed BOLD. The BOLD time-series were resampled into standard space, generating a preprocessed BOLD run in MNI152NLin2009cAsym space. First, a reference volume and its skull-stripped version were generated using a custom methodology of fMRIPrep. Several confounding time-series were calculated based on the preprocessed BOLD: framewise displacement (FD), DVARS and three region-wise global signals. FD was computed using two formulations following Power (absolute sum of relative motions)(Jenkinson et al., 2002) and Jenkinson (relative root mean square displacement between affines, Jenkinson et al.(Jenkinson & Smith, 2001)). FD and DVARS are calculated for each functional run, both using their implementations in Nipype (following the definitions by Power et al.(Jenkinson et al., 2002)). The three global signals are extracted within the CSF, the WM, and the whole-brain masks. Additionally, a set of physiological regressors were extracted to allow for component-based noise correction (CompCor)(Behzadi et al., 2007). Principal components are estimated after high-pass filtering the preprocessed BOLD time-series (using a discrete cosine filter with 128s cut-off) for the two CompCor variants: temporal (tCompCor) and anatomical (aCompCor). tCompCor components are then calculated from the top 2% variable voxels within the brain mask. For aCompCor, three probabilistic masks (CSF, WM and combined CSF+WM) are generated in anatomical space. The implementation differs from that of Behzadi et al. in that instead of eroding the masks by 2 pixels on BOLD space, the aCompCor masks are subtracted a mask of pixels that likely contain a volume fraction of GM. This mask is obtained by thresholding the corresponding partial volume map at 0.05, and it ensures components are not extracted from voxels containing a minimal fraction of GM. Finally, these masks are resampled into BOLD space and binarized by thresholding at 0.99 (as in the original implementation). Components are also calculated separately within the WM and CSF masks. For each CompCor decomposition, the k components with the largest singular values are retained, such that the retained components time series are sufficient to explain 50 percent of variance across the nuisance mask (CSF, WM, combined, or temporal). The remaining components are dropped from consideration. The head-motion estimates calculated in the correction step were also placed within the corresponding confounds file. The confound time series derived from head motion estimates and global signals were expanded with the inclusion of temporal derivatives and quadratic terms for each (Behzadi et al., 2007). Frames that exceeded a threshold of 0.5 mm FD or 1.5 standardised DVARS were annotated as motion outliers. All resamplings can be performed with a single interpolation step by composing all the pertinent transformations (i.e. head-motion transform matrices, susceptibility distortion correction when available, and co-registrations to anatomical and output spaces). Gridded (volumetric) resamplings were performed using antsApplyTransforms (ANTs), configured with Lanczos interpolation to minimize the smoothing effects of other kernels (Lanczos, 1964). Non-gridded (surface) resamplings were performed using mri_vol2surf (FreeSurfer).

Many internal operations of fMRIPrep use Nilearn 0.6.2 (RRID:SCR_001362)(Abraham et al., 2014), mostly within the functional processing workflow. For more details of the pipeline, see the section corresponding to workflows in fMRIPrep's documentation.

***Note S6.*** *The multivariate representations of conflict type and orientation are different from the congruency effect*

An explanation to the stronger encoding of conflict type in incongruent than congruent condition (Fig. 3B/D) in right 8C area may be the encoding of congruency. To test this possibility, we first tested the univariate congruency effect (incongruent minus congruent) using the parametric modulating GLM1 that was used to estimate fMRI activation levels of conflict type × congruency conditions. We observed no univariate congruency effect in the right 8C region, $t(34) = -0.03$, $p = .513$, one-tailed. Neither did we observe a multivariate congruency effect (i.e., the pattern difference between incongruent and congruent conditions compared to that within each condition) in the right 8C or any other regions. Note the definition of congruency here differed from traditional definitions (i.e., contrast between activity strength of incongruent and congruent conditions), with which we found stronger univariate activities in pre-SMA for incongruent versus congruent conditions. We further tested the possibility that the congruency effect may be manifested in behavioral relevance. To this end, we extracted the contrast of incongruent minus congruent on encoding strength of conflict similarity for each subject from the mixed-effect model based on the cross-subject RSA (see the *Representational similarity analysis* of Methods in the main text) and correlated it with the behavioral congruency effect, averaged across the five conflict types (i.e., the main effect reported in the Note S1). No significant correlation was observed ($r = 0.14$, $p = .380$, one-tailed). Taken together, these results suggested that the neural encoding strength of conflict type does not reflect the level of cognitive control engagement, but the dynamic adjustment of cognitive control instead.

Similarly, we tested whether those regions with stronger encoding of orientation in incongruent than congruent condition (i.e., right V1, V2, PF and left FEF) reflect the congruency effect. We observed no uni-voxel congruency effect in any of these regions, all uncorrected $ps > .89$, one-tailed. In addition, the orientation effect was not correlated to the behavioral congruency in any of the regions, all uncorrected $ps > .074$, one-tailed. Together with our finding that there was no correlation between the strength of orientation encoding and the conflict similarity modulation on behavioral CSEs in any of these regions (see the *Multivariate patterns of visual and oculomotor areas encode stimulus orientation* of Results in the main text), these results indicate that the encoding of orientation effect did not reflect the encoding of congruency or conflict type. Instead, we speculate that the encoding of orientations provides perceptual information to determine the conflict type.

***Note S7.*** *The cross-subject RSA captures similar effects with the within-subject RSA*

Considering the variability in voxel-level functional localizations among individuals, one may question whether the cross-subject RSA results were biased by the consistent multi-voxel patterns across subjects, distinct from the more commonly utilized within-subject RSA. We reasoned that the cross-subject RSA should have captured similar effects as the within-subject RSA if we observe the conflict similarity effect in right 8C with the latter analysis. Therefore, we tested whether the representation in right 8C held for within-subject data. Specifically, we performed similar RSA for within-subject RSMs, excluding the within-run cells. We replaced the perfectly confounded factors of conflict similarity and orientation with a common factor called *similarity_orientation*. Other confounding factor pairs (i.e., target versus response, and Stroop distractor versus Simon distractor) were addressed similarly. Results showed a significant effect of *similarity_orientation*, $t(36.3) = 3.270$, $p = .0012$, 1-tailed. Given the specific representation of conflict similarity identified by the cross-subject RSA, the within-subject data of right 8C may show similar conflict similarity modulation effects as the cross-subject data. Further research is needed to fully dissociate the representation of conflict and the representation of visual features such as orientation.

***Note S8.*** *The lateralization of conflict type representation*

We observed the right 8C but not the left 8C represented the conflict type similarity. A further test is to show if there is a lateralization. We tested several regions of the left dlPFC, including the i6-8, 8Av, 8C, p9-46v, 46, 9-46d, a9-46v (Freund, Bugg, et al., 2021). We found that none of these regions show the representation of conflict type, all uncorrected *p*s > .35. These results indicate that the conflict type is specifically represented in the right dlPFC.

***Note S9.*** *The cross-subject pattern similarity is robust against individual differences*

Due to individual differences, the multivoxel patterns extracted from the same brain mask may not reflect exactly the same brain region for each subject. To reduce the influence of individual difference, we conducted the same cross-subject RSA using data smoothed with a 6-mm FWHM Gaussian kernel. Results showed a significant conflict similarity effect, $t(1902599.9) = 5.55$, $p < .0001$, replicating the results on unsmoothed data ($t(86.8) = 5.41$, $p < .0001$).

***Note S10.*** *Cognitive control enhances target representation and suppresses distractor representation*

Using the separability of confounding factors afforded by the cross-subject RSA, we examined how representations of targets and distractors are modulated by cognitive control. The key assumption is that exerting cognitive control may enhance target representation and suppress distractor representation. We hypothesized that stimuli are represented in visual areas, so we chose a visual ROI from the main RSA results showing joint representation of target, spatial Stroop distractor and Simon distractor ($p < .005$, 1-tail, uncorrected). Only the left V4 met this criterion. We then tested representations with models similar to the main text for incongruent only trials, congruent only trials, and the incongruent – congruent contrast. The contrast model additionally used interaction between the congruency and target, Stroop distractor and Simon distractor terms. Results showed that in the incongruent condition, when we employ more cognitive control, the target representation was enhanced ($t(72.2) = 2.28$, $p = .039$, Bonferroni corrected) and both spatial Stroop ($t(85.4) = –3.94$, $p < .001$, Bonferroni corrected) and Simon ($t(38.9) = –2.80$, $p = .012$, Bonferroni corrected) distractor representations were suppressed (Fig. S8). These are consistent with the idea that the top-down control modulates the stimuli in both directions (Polk et al., 2008; Ritz & Shenhav, 2022).

### Supplementary Figures

**Fig. S1**

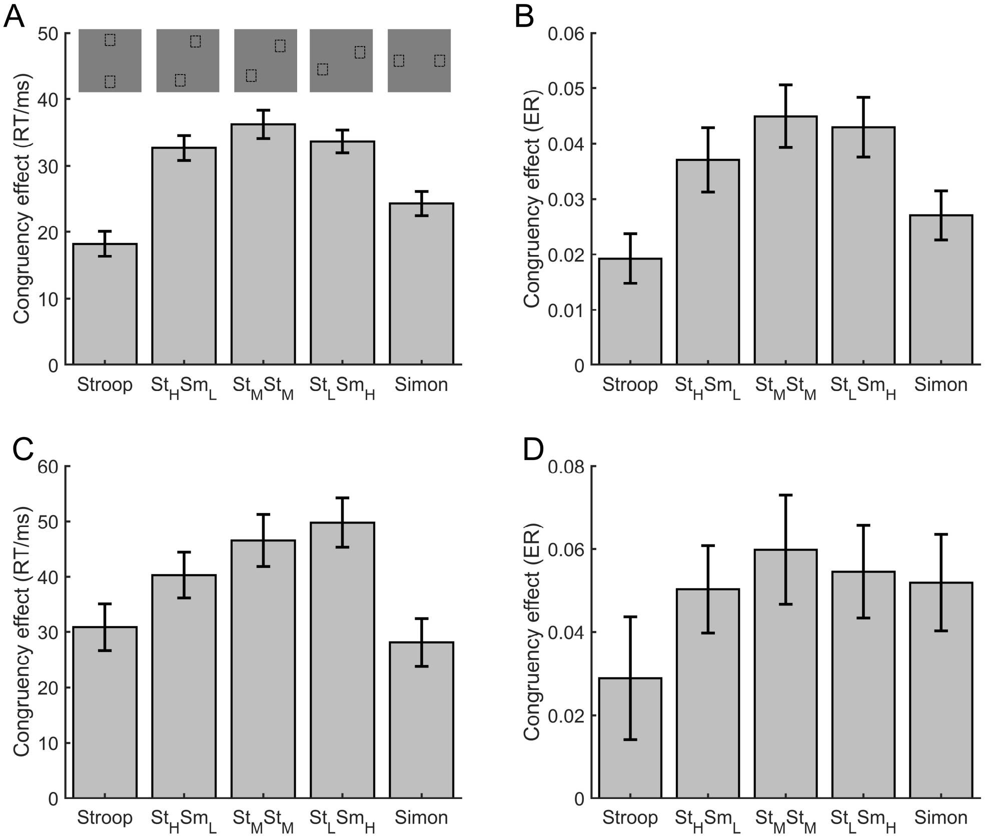


***Fig. S1.*** The congruency effects of Experiment 1 (A and B) and Experiment 2 (C and D). Error bars denote the standard errors of mean. Small insets on top of panel A denote an example of stimuli positions for each conflict type. RT = reaction time; ER = error rate.

**Fig. S2**

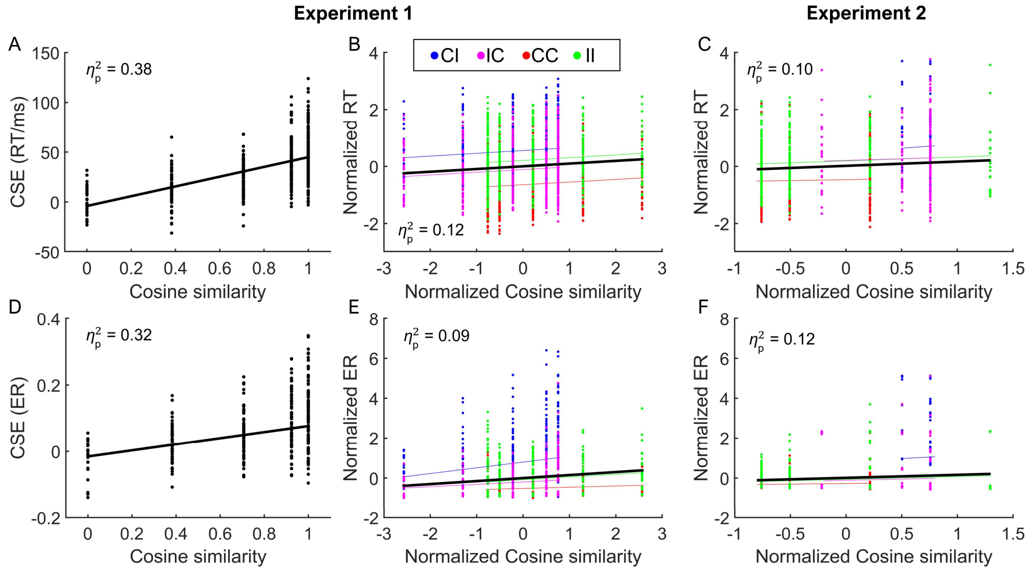


***Fig. S2.*** The conflict similarity modulation on performance of Experiment 1 (A, B, D and E) and Experiment 2 (C and F), respectively. A and D are scatter plots of CSE [i.e., (CI−CC) − (II−IC)] for RT and ER as a function of the cosine similarity, respectively. In B, C, E and F, the cosine similarity and RT / ER are normalized across conflict similarity levels within each of the four CSE conditions (i.e., CC, II, CI and IC). Conflict similarity for CC and II conditions are reversed (multiplied by −1), such that for all the four CSE conditions, higher conflict similarity is expected to be associated with worse performance (see *Behavioral analysis* in Methods). Each dot represents a subject. The thin colored lines in B, C, E and F are the fitted lines for each of the four CSE conditions, and the thick black lines are the fitted lines collapsing across all CSE conditions. For panel C and F some similarity levels are missing because of the limited trial numbers in the experimental design in Experiment 2. CSE = congruency sequence effect; RT = reaction time; ER = error rate; CI = congruent (trial n−1)-incongruent (trial n); IC = incongruent-congruent; CC = congruent-congruent; II = incongruent-incongruent.

**Fig. S3**

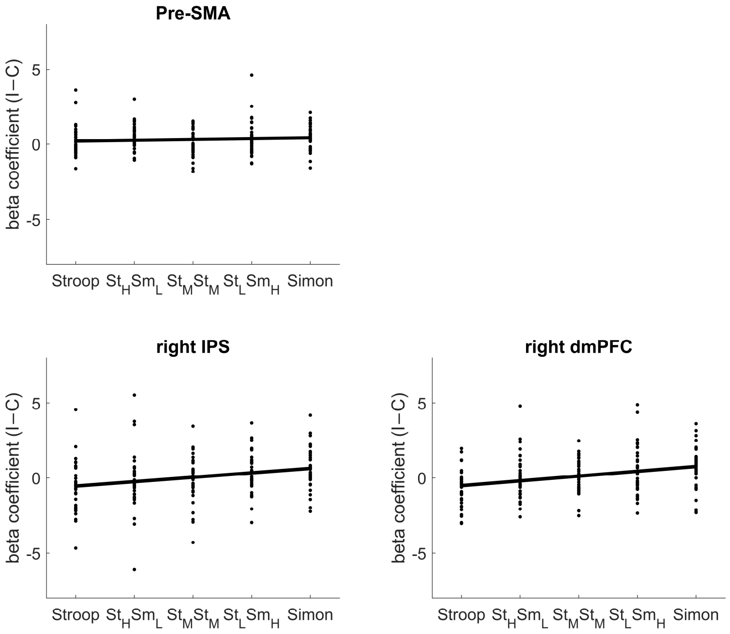


***Fig. S3.*** Neural congruency effect (I−C) by GLM2 [see the *Estimation of fMRI activity with univariate general linear model (GLM)* of Methods in the main text], plotted as a function of conflict type in different cortical ROIs. The ROIs were selected because they show a statistically significant congruency effects or parametric modulation effects when analyzed using the univariate GLM1.The pre-SMA showed overall congruency effects regardless of the conflict type (upper panel); the right IPS and right dmPFC were positively modulated by the conflict type (lower panel). Pre-SMA = pre-supplementary motor area; IPS = inferior parietal sulcus; dmPFC = dorsomedial prefrontal cortex.

**Fig. S4**

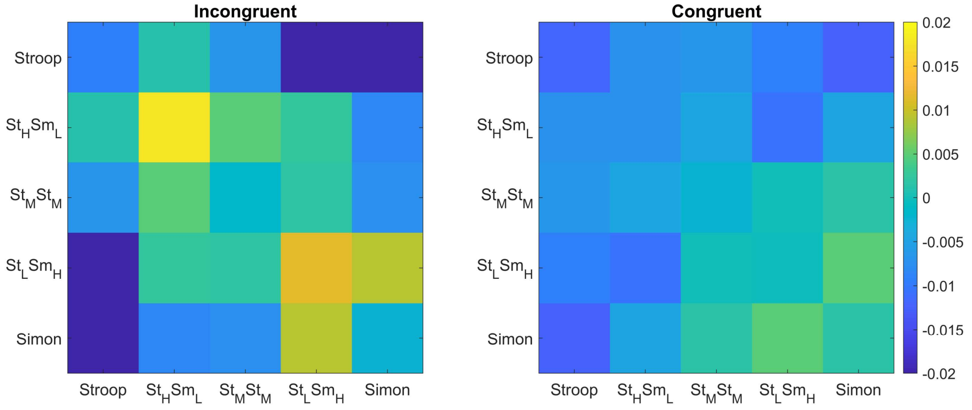


**Fig. S4.** The stronger conflict type similarity effect in incongruent versus congruent conditions. Shown are the summary representational similarity matrices for the right 8C region in incongruent (left) and congruent (right) conditions, respectively. Each cell represents the averaged Pearson correlation (after regressing out all factors except the conflict similarity) of cells with the same conflict type and congruency in the 1400×1400 matrix. Note that the seemingly disparities in the values of within-conflict cells (i.e., the diagonal) did not reach significance for either incongruent or congruent trials, $F$s $< 1$.

s

**Fig. S5**

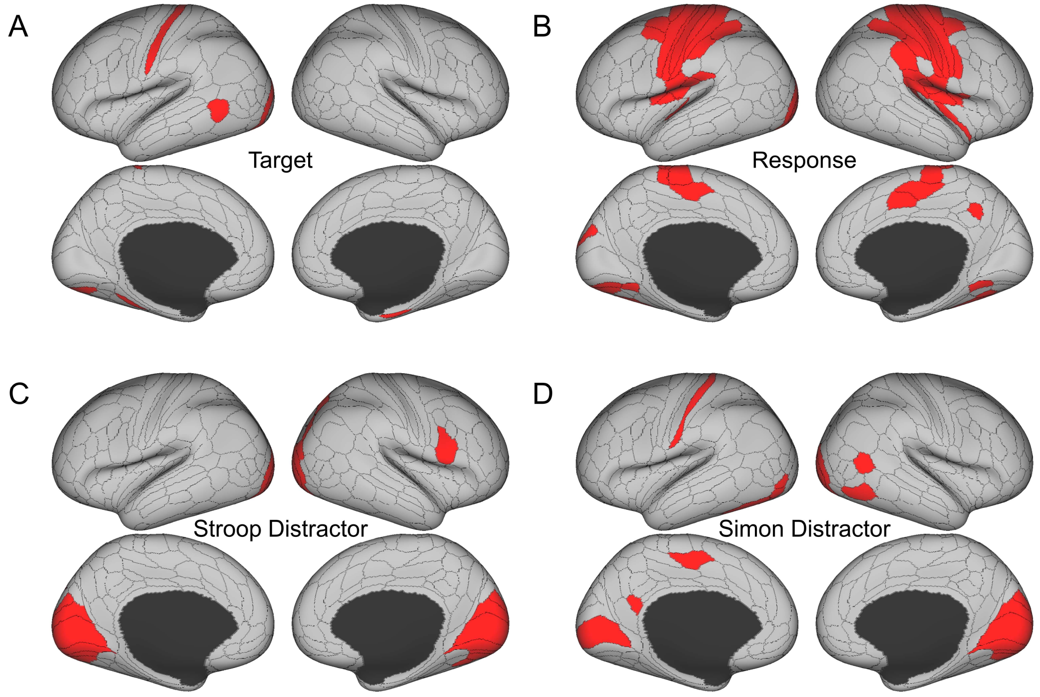


**Fig. S5.** The cortical regions showing different effects in the main RSA. (A) The target effect reflects the encoding of upward and downward arrow directions, and is mainly encoded in the visual and sensorimotor regions. (B) the response effect reflects the encoding of left and right responses, and is mainly encoded in motor regions. (C) the spatial Stroop distractor effect reflects the encoding of vertical location of the stimulus, and is encoded in bilateral visual regions. (D) the Simon distractor effect reflects the encoding of horizontal locations of the stimulus, and is mainly encoded at the bilateral visual regions. Regions in B, C and D are thresholded with Bonferroni-corrected $p < .05$ across the 360 cortical ROIs, whereas regions in A are thresholded with uncorrected $p < .005$.

**Fig. S6**

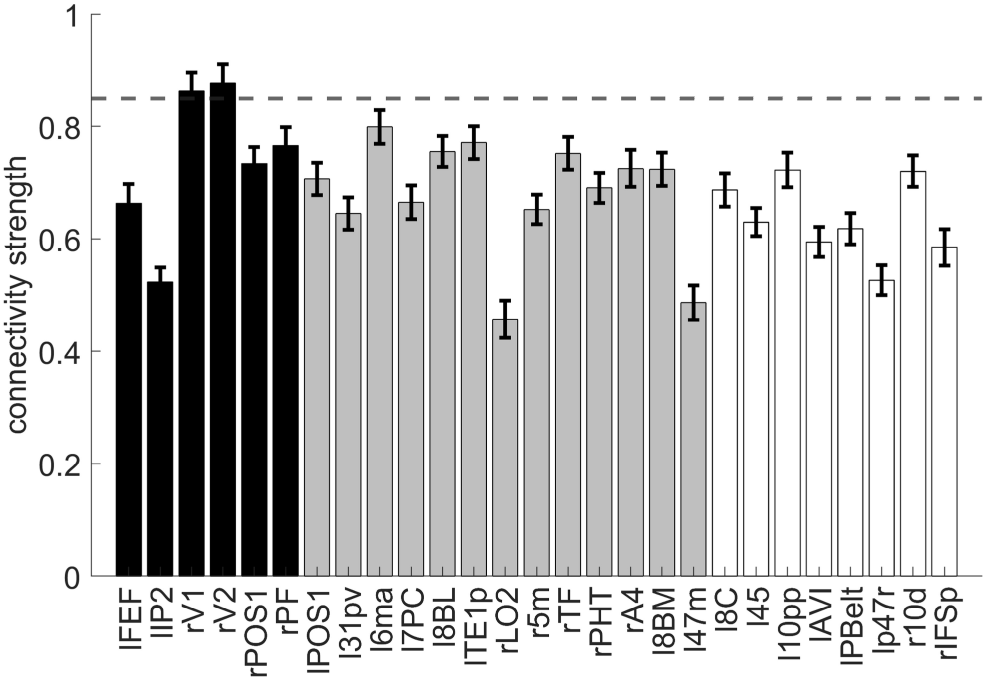


**Fig. S6.** The representational connectivity between the right 8C area and the cortical regions showing significant encoding of orientation. The black bars represent regions showing both the overall orientation effect and higher encoding strength of orientation in incongruent than congruent conditions; the grey bars are regions showing only the overall orientation effect but not higher encoding strength of orientation in incongruent than congruent conditions; and the white bars are regions not showing any of the effects of interest (i.e., uncorrected $p > 0.3$ for all the conflict type, orientation, congruency, target, response, spatial Stroop distractor and Simon distractor effects). Regions plotted in grey and white bars serve as controlled baseline. Error bars are the standard error of the mean. The dashed line indicates the upper bound of the 95% confidence interval of the highest connectivity of controlled regions (i.e., left 6ma). l = left, r = right.

**Fig. S7**

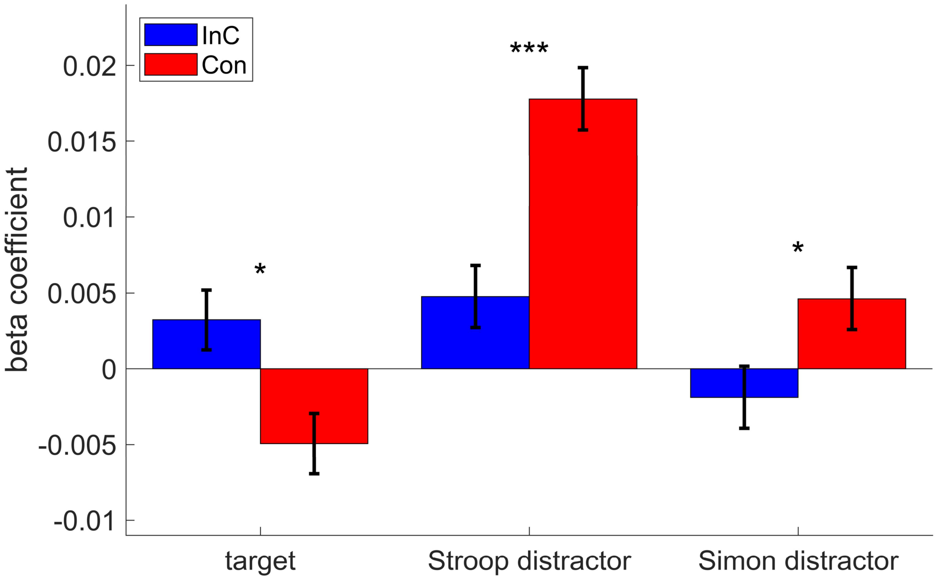


**Fig. S7.** The representational strength of target, Stroop distractor and Simon distractor in left V4 for incongruent and congruent conditions. Compared to the congruent conditions, the incongruent condition shows a stronger representation of target, but lower representation of Stroop and Simon distractors. Results are Bonferroni corrected. * $p < .05$, *** $p < .001$.

### Supplementary Tables

***Table S1.*** Brain activations for the uni-voxel parametric analysis in GLM1 (FWE corrected after probabilistic TFCE enhancement, with voxel-wise one-tailed $p < .001$ and cluster-wise $p < .05$, both one-tailed)

| Region | L/R | MNI coordinate (mm) | | | Volume (No. of voxels) | MaxZ (TFCE enhanced) | BA |
|---|---|---|---|---|---|---|---|
| | | x | y | z | | | |
| ***incongruent > congruent*** | | | | | | | |
| Pre-supplementary motor area | R | 12 | 12 | 73 | 71 | 4.18 | 6 |
| ***Positive parametric modulator (linear Simon effect)*** | | | | | | | |
| Inferior parietal sulcus | R | 52 | −64 | 33 | 81 | 4.53 | 39 |
| Dorsomedial prefrontal cortex | R | 15 | 57 | 42 | 43 | 3.92 | 9 |

Notes. L = left; R = right; BA = Brodmann area.
